# Complex three-dimensional rearing environments amplify compensatory plasticity following early blindness

**DOI:** 10.1101/2025.05.02.651847

**Authors:** Deepa L. Ramamurthy, Mackenzie Englund, Tanner J. Kovacs, Heather Dodson, Leah A. Krubitzer

**Affiliations:** Center for Neuroscience, University of California, Davis, Davis CA 95618; Department of Psychology, 900 University Ave, Riverside, CA 92521; University of Nebraska Medical Center, 42nd and Emile Streets Omaha, NE 68198

## Abstract

The neocortex has a remarkable capacity to alter its functional organization and connectivity in response to sensory loss, particularly if this loss occurs early in life. A key question is whether this cross-modal reorganization is driven by sensory deprivation or by enhanced use of the spared senses. We investigated how different rearing environments shape neural responses in primary somatosensory cortex (S1) of short-tailed opossums (*Monodelphis domestica)*, following elimination of visual inputs through bilateral enucleation in early development. Early blind and sighted littermates were reared in enriched environments to promote active tactile exploration in three-dimensional (3D) space, or in standard laboratory cages. In adulthood, both enriched groups showed adaptive changes in exploration patterns and gap crossing behaviors relative to standard-reared counterparts. Thus, early blind animals showed behavioral compensation when challenged by complex environments. Enriched rearing increased selectivity of S1 neural responses to whisker touch and altered receptive field shapes such that they were less horizontally anisotropic. This shift was strongest in enriched early blind animals, enhancing tuning along the behaviorally relevant horizontal axis more than in standard-reared early blind animals. Thus, alterations in receptive fields of neurons in S1 following early blindness were amplified by environmental complexity. Sighted opossums reared with enrichment also showed similar whisker receptive field plasticity, though to a slightly lower degree. Together, these results demonstrate the strong influence of the rearing environment on reorganization of cortex that processes inputs from the spared senses, underscoring the role of experience in directing compensatory plasticity following early sensory loss.

**SIGNIFICANCE STATEMENT:** Enhanced perceptual abilities following early sensory loss are often attributed to cross-modal recruitment of cortex linked to the deprived sense. However, plasticity also occurs within cortical areas representing spared modalities. It remains unresolved whether deprivation alone is sufficient to induce such reorganization, or whether experience using the spared sense is required. We show that enriched rearing amplifies neural coding changes in primary somatosensory cortex after early blindness, shaping receptive field geometry and promoting adaptive behavioral strategies aligned with environmental demands. Comparable changes in sighted animals reared under the same conditions reveal that reliance on touch—rather than visual deprivation alone—drives this neural and behavioral plasticity, supporting the critical role of experience in enhancing functional outcomes after early sensory impairment.

## INTRODUCTION

Early loss of vision leads to heightened perceptual abilities in the remaining senses, accompanied by structural and functional brain plasticity (Goldreich and Kanics, 2003; Wong et al., 2011; Voss, 2011; Morland et al., 2021; Mezzera and López-Bendito, 2016; Lazzouni and Lepore, 2014). However, the developmental origins of this plasticity remain poorly understood, particularly the extent to which sensory experience, movement options and affordances contribute to compensatory changes in brain and behavior. In this study, we investigate how enhancing tactile experience through rearing in an enriched environment contributes to neural and behavioral plasticity of whisker-mediated touch in a marsupial, the short-tailed opossum, when vision is lost early in development.

Exposure to enriched environments drives plasticity in sensory cortices of juvenile and adult animals with intact brains and bodies. This includes functional reorganization of sensory representations through changes in cortical territory, neuronal response properties (latency, strength, and selectivity), neuronal morphology and molecular expression (Seo, 1992; Leah et al., 1985; Coq and Xerri, 1998; Cancedda et al., 2004; Engineer et al., 2004; Polley et al., 2004; Guic et al., 2008; Mégevand et al., 2009; Devonshire et al., 2010; Alwis and Rajan, 2013, LeMessurier et al., 2018, Bibollet-Bahena 2023). Enrichment also counteracts degradation of representations after transient sensory deprivation, and promotes recovery from brain injury (Polley et al., 1999; Bartoletti et al., 2004; Sale et al, 2007; Nakashima and Dyck, 2008; Wang et al., 2013; Greifzu et al., 2014; Zhu et al., 2014; Zheng et al., 2014; Jiang et al., 2015; Kalogeraki et al., 2017; Levine et al., 2017). Rearing animals in naturalistic environments aligned with a species’ lifestyle engages ethologically relevant behavior (Landers et al., 2011; Wilkinson et al, 2010; Branchi et al., 2011; Weissbrod et al, 2013) and is especially effective in reshaping cortical sensory representations at structural, functional and molecular levels (Polley et al., 1999; Polley et al., 2004; Frostig, 2006; Gomez-Pinilla et al., 2011; Landers et al., 2011). Here, we therefore used a complex three-dimensional rearing environment designed to model aspects of the natural habitat and lifestyle of short-tailed opossums.

Many small mammals, including short-tailed opossums, routinely use forest habitats across vertical strata, from floor to canopy (Jenkins, 1974; Jones et al., 2003; Delciellos and Vieira, 2006; Abreu and Oliveira, 2014; Shapiro et al., 2014). Short-tailed opossums are terrestrial hunters and foragers (Jones et al., 2003) but prefer nesting above ground in rocky outcrops (Macrini, 2004), using branches and logs as arboreal runways to negotiate complex terrain (Lammers, 2007; Lammers and Gauntner 2008; Ladine and Kissel, 1994; Montgomery, 1980). They are skilled climbers despite lacking morphological specializations of arboreal species, relying on behavioral strategies to maintain balance and traverse arboreal substrates (Saunders et al., 1995; Saunders et al., 1998; Wilkinson et al., 2010; Schmitt et al., 2010; Rupert et al., 2014; Goin et al., 2016; Thomas et al., 2017; Lammers, 2004; Lammers, 2007; Lammers, 2009; Lammers and Biknevicius, 2004; Lammers et al., 2006; Lammers and Gauntner, 2008). Whisker touch is critical for navigating spatially complex 3D environments (Ahl, 1986; Arkley et al., 2017), guiding gap crossing and forelimb placement (Arkley et al., 2014; Niederschuh et al., 2015; Arkley et al., 2017; Grant et al., 2018). Peripheral organization of the whisker system reflects lifestyle demands and is more developed in arboreal than terrestrial species; in short-tailed opossums, whisker layout is consistent with semi-arboreal and scansorial marsupials (Beddard, 1902; Pocock, 1914; Lyne, 1958; Lyne, 1959; Muchlinski, 2013; Grant et al., 2017; Grant et al., 2018; Ramamurthy and Krubitzer, 2016; Grant et al., 2013). Thus, vertical environmental use depends heavily on behavioral modifications guided by tactile input (Lammers and Biknevicius, 2004; Lammers, 2009).

Early loss of visual inputs in short-tailed opossums at P4, prior to the formation of retinogeniculate and thalamocortical pathways, enhances tactile acuity and alters neural coding of whisker inputs in primary somatosensory cortex (S1), with receptive fields exhibiting greater selectivity along the principal axis of whisker motion (Ramamurthy & Krubitzer, 2018). Further, there is major reorganization of thalamic and cortical connections of somatosensory cortex relative to sighted animals (Dooley & Krubitzer, 2019). This raises a critical question: are alterations in somatosensory coding driven primarily by deprivation-induced anatomical reorganization, or by tactile experience itself? Here, we implement rearing conditions that encourage naturalistic tactile exploration in a spatially challenging environment designed to increase reliance on whisker use. By comparing how an enriched environment shapes neural coding and behavior related to whisker touch in sighted and early blind animals, we interrogate the role of tactile experience in directing compensatory plasticity following early blindness.

## MATERIALS AND METHODS

### Animals

Thirty-three adult short-tailed opossums of either sex (*Monodelphis domestica*; age range: 4-16 months; weight range: 61-165g) were used for electrophysiological recording experiments. Animals belonged to four experimental groups (**Table 1**), early blind and sighted reared in two conditions (standard: ‘s’, enriched: ‘e’). Previously published data from standard-reared animals were reanalyzed for comparison with enriched rearing groups (Ramamurthy and Krubitzer, 2018).

**Table 1.**
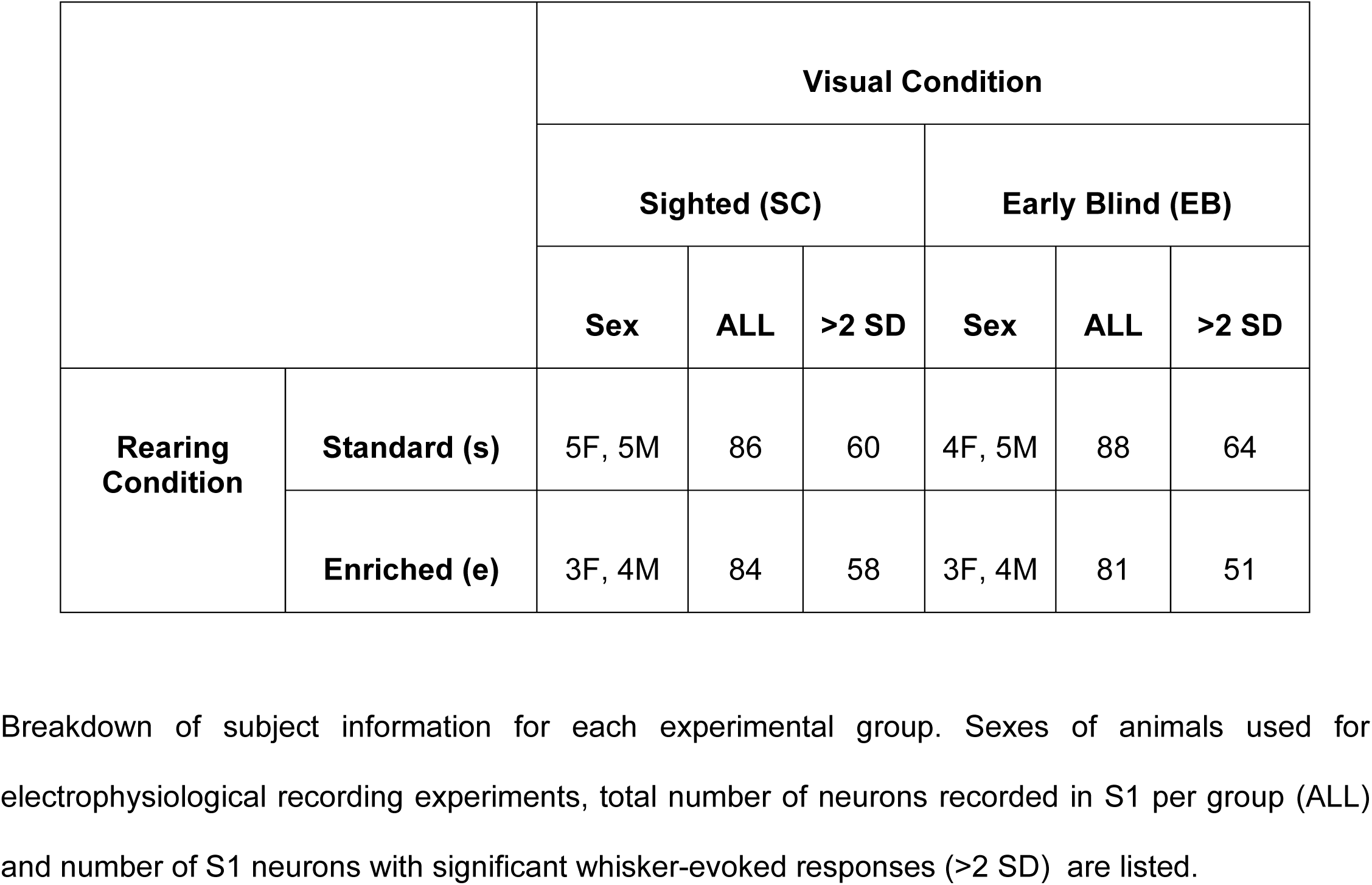
Experimental Design. Breakdown of subject information for each experimental group. Sexes of animals used for electrophysiological recording experiments, total number of neurons recorded in S1 per group (ALL) and number of S1 neurons with significant whisker-evoked responses (>2 SD) are listed.

### Ethics Statement

All protocols were approved by the Institutional Animal Care and Use Committee, and experiments were conducted according to the criteria outlined in the National Institutes of Health *Guide for the Care and Use of Laboratory Animals*.

### Bilateral enucleation

For early blind opossums, bilateral enucleations were performed at P4 (**Figure 1**). Mothers of experimental litters were lightly anesthetized with Alfaxan (initial dose: 20 mg/kg; maintenance doses: 10-50%; IM) to facilitate enucleation of the pups, which are attached to the mother at this developmental stage. Pups were anesthetized by hypothermia. Health of both the mother (respiration rate and body temperature) and the pups (heartbeat, coloration and mobility) was monitored throughout the procedure. An incision was made in the skin covering the eyes, the eyes were removed under microscopic guidance, and the skin was repositioned and resealed using surgical glue. Approximately 50% of each litter was bilaterally enucleated and the remaining littermates served as sighted controls. After complete recovery from anesthesia, mothers along with their attached litters were placed into either standard or enriched housing, depending on the experimental group.

**Figure 1.**
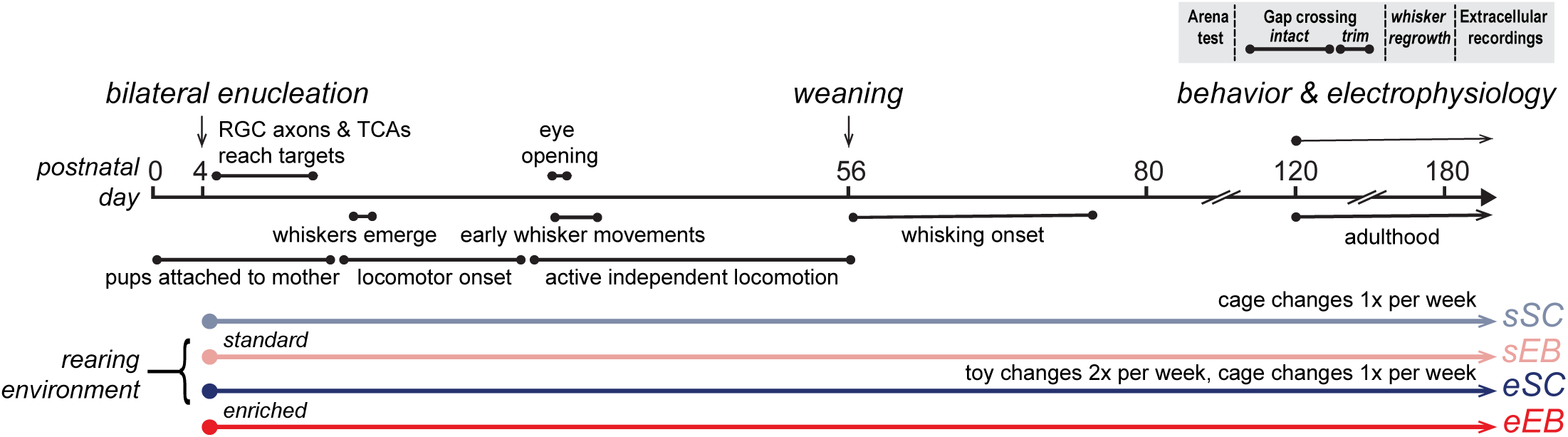
Timeline of experimental manipulations relative to developmental milestones in short-tailed opossums. The four experimental groups are indicated: sSC (standard-reared sighted control), sEB (standard-reared early blind), eSC (enriched sighted), and eEB (enriched early blind). Behavioral and electrophysiological experiments were conducted in adults following the sequence shown in the inset, but ages of adult animals were not strictly matched by postnatal day.

### Rearing paradigms

Mothers of all experimental groups were reared and housed in standard laboratory cages until bilateral enucleations were performed on the pups at P4. Following the enucleation procedure, mothers with litters belonging to the standard rearing groups were placed in standard laboratory cages, while those belonging to the enriched rearing groups were placed in custom-made enriched cages (**Figure 1-1**). All animals were maintained on a 14/10 light/dark cycle. Food and water were available *ad libitum* in the cage under both rearing conditions. All experimental litters were housed together with the mother until weaning at ∼8 weeks (P56), and at that stage they were separated from the mother, but the whole litter continued to be housed together until adulthood at ∼4 months (P120). Adults (>P120) were individually housed under standard rearing conditions, but in groups of 2-4 same-sex littermates under enriched conditions (see below).

#### Standard rearing

Under standard rearing conditions, adult animals were singly housed in standard laboratory cages, in cages similar to those typically used for laboratory rodents (cage dimensions: 18.5”l x 10”w x 8”h). The home cage was provided with a nesting cup and nesting material (shredded paper towels).

#### Enriched rearing

The enriched rearing environment in our study incorporated features common to environmental enrichment (EE) paradigms previously used in rodents (Nithianantharajah and Hannan, 2006; Sale et al., 2009; Bengoetxea et al., 2012), and additionally takes into consideration the natural lifestyle of short-tailed opossums (**Figure 1-1**). Juvenile animals were cohoused until weaning with the mother as well as littermates, and then with just their littermates from weaning until adulthood, as described above. Enriched cages were provided with standard bedding and nesting material, as well as some additional nesting material with a different texture. In addition to the nesting cup placed on the floor as in standard cages, enriched cages provided the option of nesting high above the cage floor, in nesting boxes attached to the cage walls. A single nesting box was provided in the cage at juvenile stages of the experimental litters. In adulthood, balanced numbers of eSC and eEB littermates were housed together in each enriched cage, with a maximum of four opossums per cage. Each adult animal was provided with an individual nesting box. Unlike common enrichment paradigms, which typically incorporate toys into standard cages, the enriched cages used here were substantially larger (∼8x in volume; 29”l x 18”w x 24”h) compared to standard cages to encourage tactile exploration in all three dimensions. The greatest increase relative to standard cages was along the vertical dimension: enriched cages were 3x taller than standard cages. Enrichment toys (Bio-Serv) of different shapes and textures were provided in the cage. Toys were replaced, and the positions of manzanita branches were changed every 2–3 days to introduce novelty and promote continual exploration of the environment.

### Behavioral testing

#### Arena testing

Arena testing was performed in three adult animals from each of the four experimental groups. The arena consisted of a cage with the dimensions of enriched cages, containing a single nest box near the top of the arena, with a branch leading up to it. Animals were placed in the arena and allowed to explore it for 5 minutes in complete darkness, and movies of their behavior were acquired under infrared (IR) illumination. idTracker software (**Figure 2-1A**; Romero-Ferrero et al., 2019) was used to extract movement trajectories of individual animals, which was further analyzed in MATLAB. In each movie frame, we segmented the arena into a vertical zone (cage walls, branches, and nest area) and a horizontal zone (cage floor). Exploratory activity was quantified from session-wise binarized matrices (1280 × 720 pixels; height × width) containing the animal’s position in each frame. To assess differences in how animals sampled vertical space in the arena, binarized occupancy maps were summed across sessions separately for standard- and enriched-reared animals and collapsed along the horizontal axis, yielding a per-row estimate of vertical occupancy. Vertical occupancy profiles were then binned in 20-row increments (20 px per bin; vertical dimension only) and normalized to the percentage of total vertical occupancy (summing to 1 across the vertical axis). For each animal, total vertical occupancy (V) and horizontal occupancy (H) were computed by summing binarized pixel values within the corresponding zone across all frames. A normalized vertical preference index was calculated for each animal as:

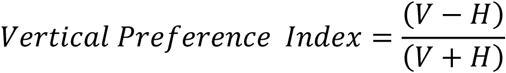

Additional analysis of exploratory behavior in the arena test was performed through manual scoring in BORIS (Behavioral Observation Research Interactive) software. All home cage behavior data were analyzed offline by four independent observers such that multiple epochs were scored by at least two independent observers with a high interrater reliability (Cohen’s κ coefficient of > 0.85; κ = 0 indicates no agreement in scoring between observers and κ = 1 indicates perfect agreement; Ramamurthy & Krubitzer, 2018; Landis and Koch, 1977). Arena exploration was initially scored using a binary scale of active states (locomotion bouts) and inactive states (resting bouts). A switch between an active and inactive state was only considered to have occurred if a bout lasted for at least 5 s. Active states were then further scored for crossing events that occurred during natural exploration of the arena. Crossing events were defined as instances in which an animal crossed from one surface to another within the arena. Four types of crossing events were scored: (1) floor to wall/wall to floor (2) wall to wall (3) floor to branch/branch to floor (4) nest or branch to wall/wall or branch to nest. Since systematic differences were not observed across crossing event types, only combined summary plots are shown.

#### Gap crossing task

Gap crossing was tested in 3-4 adult animals per experimental group. Animals were allowed to spontaneously cross from one platform (“start”) to another platform (“goal”) with either small (2-10 cm) or large gap distances (12-14 cm) separating them, which were interleaved in a pseudorandom order. Animals were tested for 7 days with intact whiskers, followed by 1-3 days of testing after whisker trimming. On each trial animals were allowed to remain on the start platform and attempt crossing for a maximum of 1 minute. If no crossings were performed, the trial was ended, and animals were removed from the platform for 1 min before returning to the platform for the next trial. Movies of this behavior were acquired under IR illumination or dim red light, and DeepLabCut was used for analysis of gap crossings (**Figure 2-1B-C**). The snout tip, the base of the tail and the position of the edges of each platform were labeled in a subset of frames to generate the dataset (260 frames across 13 opossums) on which the network was trained (200,000 iterations). On a subset of trials, slow-motion movie clips in a zoomed-in field of view were acquired to visually confirm the use of whiskers during gap crossing. Infrared motion sensors (emitter/detector pairs) placed on either side of the edge of each platform detected beam breaks and beam break times were recorded as a secondary method of confirmation.

### Surgical procedures

Animals were anesthetized with urethane (initial dose: 1.25 g/kg, 30% in saline, IP; supplemental doses: 0.125-0.313 g/kg, 30% in saline, IP). Vital signs (respiration and body temperature) were monitored throughout the experiment. At the beginning of the surgery, dexamethasone (0.4-2.0 mg/kg; IM) was administered to minimize intracranial swelling. Lidocaine (2% solution; SC) was injected at the midline of the scalp and around the ears. The animal was then placed in a stereotaxic frame. Following an incision at the midline of the scalp and retraction of the temporal muscle, a craniotomy was performed to expose the entire parietal cortex and the dura was retracted. The surface of the neocortex was covered with a layer of silicone fluid to prevent desiccation and then photographed so that electrode penetration sites could be related to patterns of vasculature. After the craniotomy, the animal was removed from the stereotaxic frame to permit full access to the face for whisker stimulation. Three stainless steel skull screws were inserted in the skull contralateral to the recording electrode. The head of the animal was stabilized by cementing a head post to the skull screw in a rostral location over the cortex. The other two skull screws were inserted epidurally into the skull over the cerebellum and over the olfactory bulbs, serving as the reference and ground electrodes, respectively, and also providing additional anchoring support for the dental cement.

### Extracellular recordings and whisker stimulation

Somatosensory receptive fields were first coarsely mapped by recording multiunit activity while applying tactile stimuli using a handheld probe. The representation of the whiskers in S1 was localized by progressively testing electrode penetration sites medial to the rhinarium representation and caudal to the lower jaw/lip representation. After this, when receptive fields were found to be located on the mystacial or genal whisker pad (Ramamurthy & Krubitzer, 2016), computer-controlled whisker deflections were used to quantitatively measure single unit receptive fields.

Whisker stimuli consisted of single whisker deflections (2°) delivered using piezoelectric actuators (4 ms/100 ms/4 ms ramp-hold-return; 5 mm from the whisker base; 1.4 mm forward excursion). Data were collected for a grid of sixteen whiskers on the mystacial pad to enable construction of somatotopic receptive fields. Whiskers were deflected in a pseudorandom order with 50-100 trials collected per whisker (in blocks of 3-50 trials) at each recording site. Extracellular recordings (amplifier gain: 10,000X; A-M Systems Model 1800 Microelectrode AC Amplifier; A-M Systems, Carlsborg, WA; sampled at 28 kHz; Power1401, Cambridge Electronic Design Limited, Cambridge, UK) were made 400-500 µm below the pial surface using monopolar tungsten microelectrodes (FHC, Inc., Bowdoin, ME; 1-5 MΩ at 1 kHz) lowered with a hydraulic microdrive (David Kopf Instruments, Tujunga, CA). Recordings were bandpass filtered (300-3000 Hz), and spike-sorting was carried out offline to confirm isolation of single units. After electrophysiological data was collected, fluorescent probes were used to mark the location of recording sites, so that these could be related to histologically identified cortical field boundaries (**Figure 3-1**).

### Histology

At the end of each experiment, animals were injected with an overdose of sodium pentobarbital (Beuthanasia; 250 mg/kg IP) and transcardially perfused with 0.9% saline, followed by 2-4% paraformaldehyde in phosphate buffer, and then 2-4% paraformaldehyde in 10% phosphate-buffered sucrose. The brain was extracted and post-fixed (1-2 hours in 4% paraformaldehyde in 10% phosphate-buffered sucrose). In some cases, the cortical hemispheres were flattened and the tissue was sectioned tangentially at 30 μm using a freezing microtome, and then processed for myelin staining (Gallyas, 1979; Dooley et al, 2013) to enable reconstruction of cortical field boundaries. Reconstructions were drawn using Adobe Illustrator CS5 (Adobe, San Jose, CA). Digital images of processed tissue were obtained using either a Nikon Multiphot system or an Optronics MicroFire digital microscope camera. Contrast and brightness of whole images were adjusted using Adobe Photoshop CS5 (Adobe, San Jose, CA).

### Neural data analysis

Data files were processed blind to experimental group identity. Single-unit isolation was performed while running the experiment using template-matching procedures in Spike2 (Cambridge Electronic Design Limited, Cambridge, UK; RRID:SCR_000903), and unit isolation was verified offline in Spike2, using principal component analysis. Stable waveforms and firing rates over the course of the recording session, and <0.5% refractory period violations (interspike interval <1.5 ms) were required for units to be included in the analysis. Spike times were then exported to MATLAB (Mathworks, Inc., Natick, MA; RRID:SCR_001622) and analyzed using custom scripts. Whisker evoked firing rates were measured as the spike count following stimulus onset (0-100 ms) and were considered significant if they exceeded prestimulus spike counts in an equivalent time window by two standard deviations. The *best whisker* (BW) was defined as the whisker with the highest evoked response, and depending on their proximity to the BW the remaining whiskers were categorized as *first-order surround whiskers* (1*°* SW; immediately adjacent to BW), *second-order surround whiskers* (2*°* SW; one position removed from BW) and *third-order surround whiskers* (3*°* SW; two positions removed from BW).

One measure of receptive field size used was the number of whiskers that evoked a significant response in each recorded neuron. Another measure used was the rank-ordered whisker tuning curve for each neuron, where responses to all whiskers tested were sorted in descending order of whisker-evoked response magnitude, and normalized to peak (BW response). Further, mean 2D somatotopic receptive fields were obtained after normalizing SW-evoked response magnitudes to the BW-evoked response, and aligning receptive fields of single units such that the BW was at the center of the receptive field for each unit. Because *Monodelphis* lacks discrete anatomical cortical barrels (Ramamurthy & Krubitzer, 2016), 2D receptive fields were aligned relative to the BW position rather than an anatomically defined “principal whisker” or “columnar whisker”. 50, 75, and 90% contour lines (for response levels relative to the BW) were plotted for the mean receptive fields (smoothed by linear interpolation) to facilitate visualization. Raw values were used for all calculations and statistical comparisons.

We define anisotropy in receptive fields as the difference in the tuning width of an S1 neuron’s receptive field when measured along different spatial directions of the whisker pad—across arcs (horizontal axis) and across rows (vertical axis). Anisotropies in the structure of somatotopic receptive fields were quantified as the *shape index:*

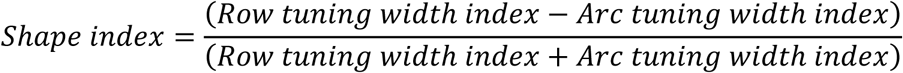

If the sum of the row and arc tuning width indices was zero, the shape index was assigned to be zero. Shape indices ranged from −1 to 1, with values >0 indicating broader tuning along the row compared to the arc (i.e. broader horizontal tuning), and values <0 indicating broader tuning along the arc compared to the row (i.e. broader vertical tuning). Neuronal selectivity for the best whisker along the row and arc dimensions was measured by assessing tuning width separately along the horizontal and vertical axes as the *row tuning width index* and the *arc tuning width index* respectively, either considering only 1*°* SW, or 1*°,* 2*°*, 3*°* SW in the row or arc of the BW, as stated below:

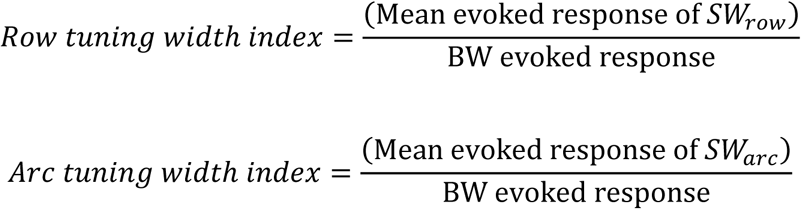

Additionally, the *whisker selectivity index (WSI)* for BW vs. in-row or in-arc whiskers was computed separately for each neuron, considering BW responses relative to 1*°,* 2*°*, 3*°* SW responses.

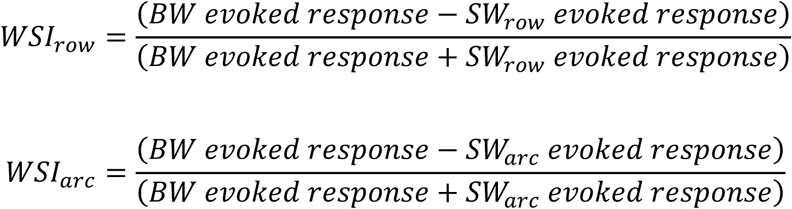

In-row and in-arc selectivity indices ranged from 0 to 1, with values closer to 1 indicating higher selectivity for the BW in the corresponding dimension.

Receptive field shape was quantified at two spatial scales (1° only vs. 1°-3°) because somatotopic receptive fields of neurons in the S1 whisker representation consist of the BW (peak response) and a graded, multi-whisker surround shaped by intracortical integration, including anisotropies along row vs. arc axes. This receptive field organization has been demonstrated in classical mouse and rat studies (e.g., Simons, 1978; Brecht & Sakmann, 2002; Fox, 2008), as well as prior work demonstrating similar receptive field structure in opossum S1 (Ramamurthy & Krubitzer, 2016; Ramamurthy & Krubitzer, 2018).

Since receptive field shape analysis characterizes the overall geometry of the field around the BW, it cannot be meaningfully computed for a non-continuous portion of the surround alone (e.g., only 2°/3° SWs); therefore 2°/3° were not analyzed separately from 1° SWs; instead, shape analyses with only 1° SWs characterized local receptive field configuration, whereas analyses including 1°-3° SWs captured receptive field configuration shaped by broader spatial integration.

### Statistical Analysis

Statistical analyses were performed in MATLAB and R. Permutation tests for the difference in medians were used to assess differences between two groups (referred to in brief as ‘permutation test’). Fisher’s exact tests (two-sided) were used for analyses of 2×2 contingency tables to compare proportions across conditions. All statistical tests for neuronal data use ‘n’ of cells. We accounted for interindividual variability using linear mixed effects models.

To model crossing behavior in the gap-crossing task, we fit a generalized linear mixed-effects model using ‘fitglme’ in MATLAB (Poisson distribution; log link). Crossing counts were modeled with the log of observation time included as an offset to estimate rates. Fixed effects included experimental group, gap size, whisker trimming, and the interactions of experimental group with gap size and with whisker trimming, respectively. Since whisker trimming abolished crossings on 100% of large-gap trials, those trials were excluded from this analysis. Animal identity was included as a random intercept (1|ID) to account for repeated measures. The model specification was as follows:

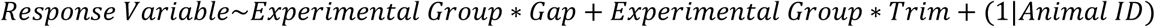

The model was estimated using the Laplace approximation for maximum likelihood. Overdispersion was evaluated via Pearson dispersion; if dispersion exceeded 1.5, we added an observation-level random effect (1|Obs) to handle overdispersion.

To analyze gap crossing durations on successful trials we fit a linear mixed-effects model using ‘fitlme’ in MATLAB with the specification:

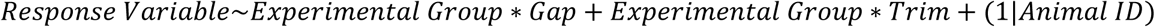

where experimental group and test condition were modeled as fixed effects, and animal ID modeled as a random effect.

Planned post hoc comparisons for both models were performed using coefTest with contrast matrices for group and condition effects (gap size and whisker trimming), including a difference-in-differences contrast for the blindness × environment interaction. ‘coefTest’ returned p-values for each contrast, with Satterthwaite degrees of freedom applied where applicable.

For neural data, experimental group was modeled as a fixed effect and animal identity as a random effect, using the model specification:

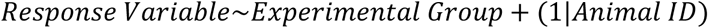

Planned contrasts were performed using ‘coefTest’ for pairwise group comparisons and for a difference-in-differences (blindness × environment) contrast, with p-values returned and Satterthwaite degrees of freedom applied where applicable. Additional models were fit for key analyses with sex included as a fixed effect to test for differences between males and females, but no significant sex differences were identified.

Exact p-values for fixed effects of experimental group and/or test condition (gap size/whisker trimming) obtained from Poisson and Gaussian generalized linear mixed-effects models, or from Fisher’s exact tests for categorical comparisons, with pairwise contrasts across experimental groups are provided in **Table 2** and **Table 3**.

**Table 2.** Statistical comparisons for key behavioral results. Exact p-values for fixed effects of experimental group and/or test condition obtained from Poisson and Gaussian generalized linear mixed-effects models, or from Fisher’s exact tests for categorical comparisons, with pairwise contrasts across experimental groups.

| Result/<br>Figure Panel | Fixed<br>Effect<br>p-value | Pairwise comparisons<br>p-values |  |  |  |  |  |
| --- | --- | --- | --- | --- | --- | --- | --- |
|  | Test<br>Condition | sSC | sEB | eSC | eEB |  |  |
| % successful trials<br>**Fig. 2B**<br>Alpha with Bonferroni<br>correction<br>= 0.0125<br>(1) large vs. small<br>(2) intact vs. trim | - | - | - | - | - |  |  |
|  |  | <2.2e-16 | <2.2e-16 | <2.2e-16 | <2.2e-16 |  |  |
| # Gap crossings/minute<br>**Fig. 3C**<br>(1) large vs. small<br>(2) intact vs. trim | 2.4277e-09 | 2.4277e-09 | 1.7407e-09 | 9.8418e-10 | 2.0062e-09 |  |  |
|  | 0.5356 | 0.5356 | 0.37091 | 0.42035 | 0.15936 |  |  |
| Gap crossing duration<br>**Fig. 3D**<br>(1) large vs. small<br>(2) intact vs. trim | 1.1011e-79 | 8.0737e-92 | 1.4663e-08 | 0.0128 | 0.4872 |  |  |
|  | 2.888e-07 | 0.0081 | 9.6063e-09 | 7.2433e-05 | 0.1481 |  |  |
|  | **Expt.**<br>**Group** | **sSC vs.sEB** | **sSC vs. eSC** | **sSC vs. eEB** | **sEB vs. eEB** | **sEB vs. eSC** | **eSC vs. eEB** |
| # Gap crossings/minute<br>**Fig. 3C**<br>(3) across groups<br>(large, intact) | 0.5927 | - | - | - | - | - | - |
| # Gap crossings/minute<br>**Fig. 3C**<br>(3) across groups<br>(small, intact) |  | - | - | - | - | - | - |
| # Gap crossings/minute<br>**Fig. 3C**<br>(3) across groups<br>(small, trim) |  | - | - | - | - | - | - |
| Gap crossing duration<br>**Fig. 3D**<br>(3) across groups<br>(large, intact) | 6.8796e-51 | 7.7220e-35 | 3.6476e-62 | 3.3333e-72 | 3.2002e-09 | 3.8990e-04 | 0.0021 |
| Gap crossing duration<br>**Fig. 3D**<br>(3) across groups<br>(small, intact) |  | 0.7771 | 0.0691 | 2.9099e-14 | 1.0844e-13 | 0.1031 | 7.5517e-05 |
| Gap crossing duration<br>**Fig. 3D**<br>(3) across groups<br>(small, trim) |  | 1.3303e-05 | 5.6083e-19 | 6.8960e-15 | 6.6591e-04 | 8.6167e-06 | 0.3183 |

Exact p-values were obtained from Poisson and Gaussian generalized linear mixed-effects models, or from Fisher's exact tests for categorical comparisons, with pairwise contrasts across experimental groups as indicated (see Methods).

**Table 3.** Statistical comparisons for key electrophysiological results. Exact p-values for fixed effects of experimental group from Gaussian generalized linear mixed-effects models, with pairwise contrasts across experimental groups.

| Result/<br>Figure Panel | Fixed<br>Effect<br>p-value | Pairwise comparisons<br>p-values |  |  |  |  |  |
| --- | --- | --- | --- | --- | --- | --- | --- |
|  |  | sSC vs.sEB | sSC vs. eSC | sSC vs. eEB | sEB vs. eEB | sEB vs. eSC | eSC vs. eEB |
| Spontaneous activity<br>**Fig. 3C** | 0.856 | - | - | - | - | - | - |
| Whisker-evoked activity<br>**Fig. 3D** | 4.68e-10 | 4.53e-04 | 4.09e-04 | 1.11e-11 | 0.939 | 1.38e-04 | 3.04e-04 |
| RF size: # of whiskers<br>**Fig. 3E (left)** | 1.58-07 | 9.43e-07 | 4.29e-06 | 8.86e-07 | 0.832 | 0.763 | 0.621 |
| Rank-ordered whisker tuning curve<br>**Fig. 3E (right)** | 1.55e-25 | 3.271e-11 | 1.88e-22 | 4.04e-19 | 8.19e-04 | 0.006 | 0.618 |
| Shape index: 1<sup>st</sup> order adjacent whiskers<br>**Fig. 4C (left)** | 0.004 | 0.249 | 0.445 | 6.84e-04 | 0.628 | 0.023 | 0.0033 |
| % horizontal: 1<sup>st</sup> order adjacent whiskers<br>**Fig. 4C**<br>Alpha with Bonferroni correction = 0.008 (right) | - | 0.266 | 0.185 | 0.001 | 0.039 | 0.855 | 0.083 |
| Shape index: 1<sup>st</sup> – 3<sup>rd</sup> order adjacent whiskers<br>**Fig. 4D (left)** | 3.45e-05 | 0.093 | 0.013 | 2.19e-06 | 0.384 | 0.001 | 0.016 |
| % horizontal: 1<sup>st</sup> – 3<sup>rd</sup> order adjacent whiskers<br>**Fig. 4D**<br>Alpha with Bonferroni correction = 0.008 (right) | - | 0.516 | 9.88e-05 | 2.00e-06 | 2.50e-05 | 7.73e-04 | 0.340 |
| % BW response: in-arc whiskers<br>**Fig. 4E (left)** | 0.042 | 0.0325 | 0.0116 | 0.0244 | 0.652 | 0.812 | 0.848 |
| % BW response: in-row whiskers<br>**Fig. 4E (left)** | 8.51e-20 | 7.10e-05 | 7.56e-14 | 1.12e-18 | 6.19e-05 | 1.25e-08 | 0.056 |
| BW vs. in-arc whiskers<br>**Fig. 4F (left)** | 0.043 | 0.080 | 0.014 | 0.015 | 0.436 | 0.419 | 0.957 |
| BW vs. in-row whiskers<br>**Fig. 4F (right)** | 2.49e-18 | 2.86e-04 | 8.18e-12 | 7.00e-18 | 3.03e-04 | 8.31e-09 | 0.018 |

Exact p-values were obtained from Gaussian generalized linear mixed-effects models, with pairwise contrasts across experimental groups as indicated (see Methods).

## RESULTS

### Behavioral adaptations of blind and sighted opossums in enriched rearing environments

In short-tailed opossums, postnatal day 0 (P0) is developmentally equivalent to embryonic stages in rodents (approximately E11 in mouse and E12 in rat; Molnár and Blakemore, 1995; Molnár et al., 1998). Bilateral enucleations were conducted at P4, before retinal afferents and thalamocortical axons reached their respective targets (Taylor and Guillery, 1994; Molnár et al., 1998), completely eliminating both spontaneous retinal activity and visually evoked input (**Figure 1)**. Following bilateral enucleation on postnatal day 4, opossum mothers and their litters, including sighted and blind pups, were transferred into a new standard or enriched home cage environment (**Figure 1**; **Figure 1-1A-D**). Prior to P12, sensory experience in either environment was determined entirely by behavioral patterns of the mother, due to the obligate attachment of opossum pups. Juvenile animals continue to rely on the mother for both sustenance and transportation until weaning at P56, but with increasing independent exploration (**Figure 1-1E-F**). Opossum mothers always preferentially nested high above the ground (n = 3 mothers, producing a total of 4 enriched experimental litters; example in **Movie 1**), so pups were forced to rely heavily on touch to navigate the enriched environment. By adulthood, both sighted and early blind animals were highly adapted to their environments (**Movie 2-3**).

To assess how rearing condition and visual condition (blind versus sighted) shape exploratory behavior, we tested each group in a novel arena matching the dimensions and basic layout of the enriched home cages (**Figure 2; Figure 2-1A**). We quantified spatial sampling preferences (**Figure 2A-C** and **2E**), overall locomotor activity (**Figure 2D** and **2F**), and spontaneous crossings between surfaces during natural exploration (**Figure 2D** and **2G**), capturing both spatial and temporal aspects of behavior. These measures contextualize and complement the more controlled gap crossing behavior assay presented in the next section. Movement trajectories were highly similar for sighted and blind animals within each rearing condition (**Figure 2A-B**; **Figure 2-1A**), consistent with successful behavioral compensation by blind animals even in more spatially complex environments. In contrast, rearing environment strongly impacted how animals sampled the novel arena: enriched opossums explored vertical space more extensively than those reared in standard cages (p = 0.018, permutation test; **Figure 2C**; **Figure 2E**), though overall activity levels (p=0.565, permutation test) and durations of naturally occurring crossing events were not significantly different (p=0.232, permutation test; **Figure 2D**; **Figure 2F-G**). Quantifying gap crossing during free exploration has some limitations which may mask differences in sensory-mediated behavior. Crossing events that occurred during free exploration, where animals crossed from surface to surface did not necessarily involve crossing of discrete gaps, and we could not control for gap size across different crossing events. Further animals were free to adopt varying postures prior to crossing, and to use varying behavioral strategies involving sensory inputs from multiple body parts to aid gap crossing during free exploration.

**Figure 2.**
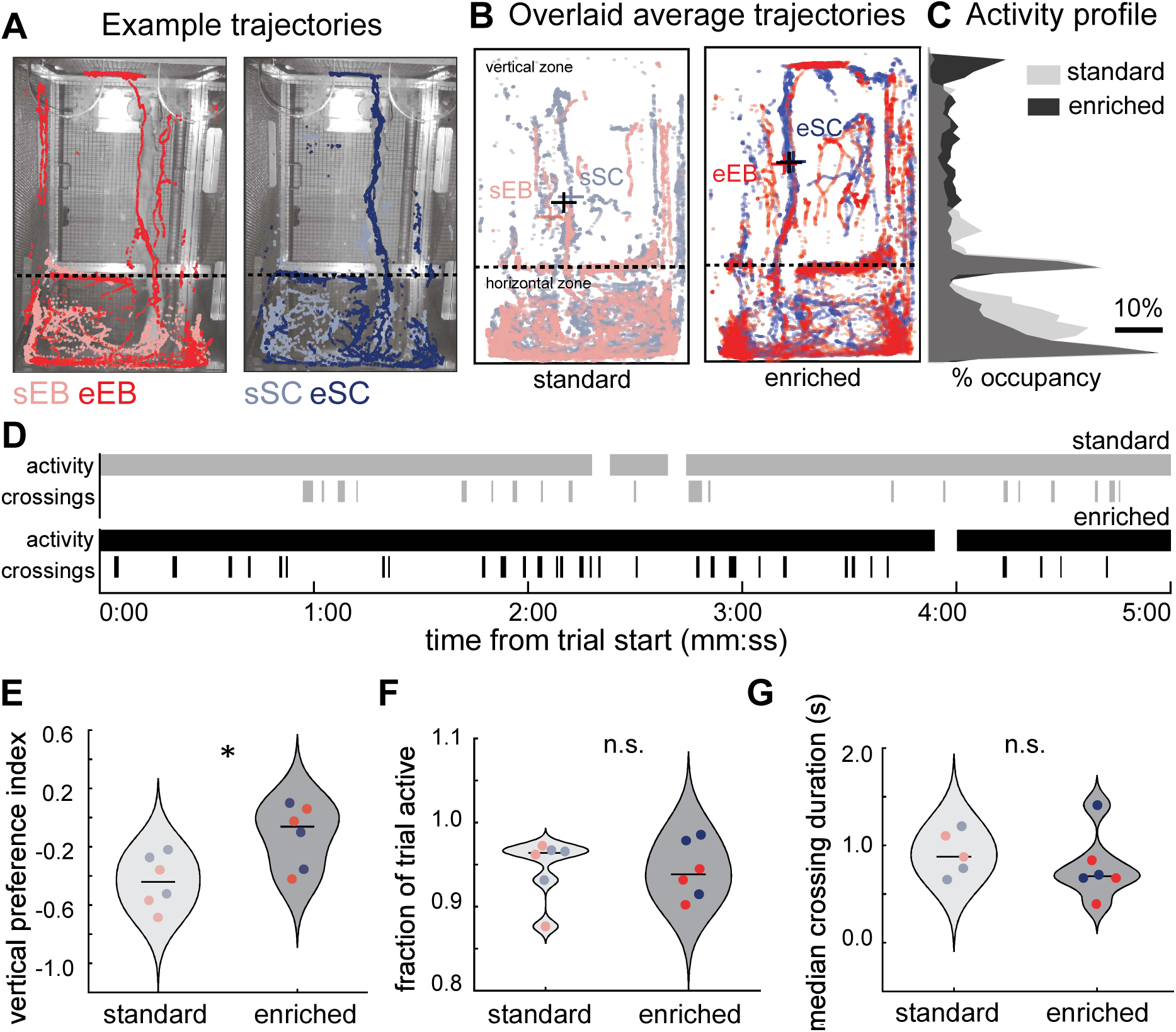
Enriched rearing alters spatial patterns of exploratory behavior. **A.** Example movement trajectories from one animal in each experimental group (sEB: light red, eEB: dark red, sSC: light blue, eSC: dark blue) tracked over five minutes during behavioral testing in an enriched arena. **B.** Overlaid movement trajectories from all animals in each experimental group in the arena test show a pattern of increased exploration of vertical vs. horizontal space in opossums reared in enriched cages compared to those reared in standard cages. Plus symbols indicate the center of mass of movement trajectories for each group (colors as in **A**), with data for animals combined within each rearing condition shown in black. **C.** Vertical occupancy profiles during arena exploration. Normalized distributions of vertical position (height above the arena floor) during exploration derived from 1280×720 px video frames (20 px bins used for visualization), pooled across animals (black: enriched, gray: standard; dark gray denotes regions of overlap between the two distributions. **D.** Ethograms showing the temporal structure of locomotor activity and spontaneous crossing events during arena exploration. Horizontal bars indicate periods of activity for representative trials from standard-reared (top, gray) and enriched-reared (bottom, black) animals. Vertical tick marks denote spontaneous crossings between surfaces. Time is plotted relative to trial start (mm:ss). **E-G**. Summary behavioral metrics during arena exploration in standard and enriched animals. Distributions of vertical preference index (**E**), fraction of trial time spent active (**F**), and median crossing duration (**G**). Violin plots show Gaussian kernel density estimates of the pooled distribution for each category; horizontal lines denote medians. Points represent individual animals, color-coded by experimental group. Animals reared in the enriched environment show significantly a signficant increase in vertical prefernce index. Overall activity levels and duration of crossing events during free exploration were not signficiantly different.

To assess sensorimotor performance under more precisely defined task parameters, we used a gap crossing task (conducted in darkness) in which gap size was systematically varied across trials, animals had only one possible path forward (across the gap), and the constraints of the apparatus required animals to adopt a standardized posture prior to gap crossing (**Figure 3A**; **Figure 2-1B-C**). Crossing large gaps required whisker use, as the goal platform was out of reach from the start platform except through whisker contact. We quantified (1) the fraction of trials resulting in a successful crossing (**Figure 3B**), (2) crossing rates over time (**Figure 3C**), and (3) the duration of successful crossing events (**Figure 3D**).

**Figure 3.**
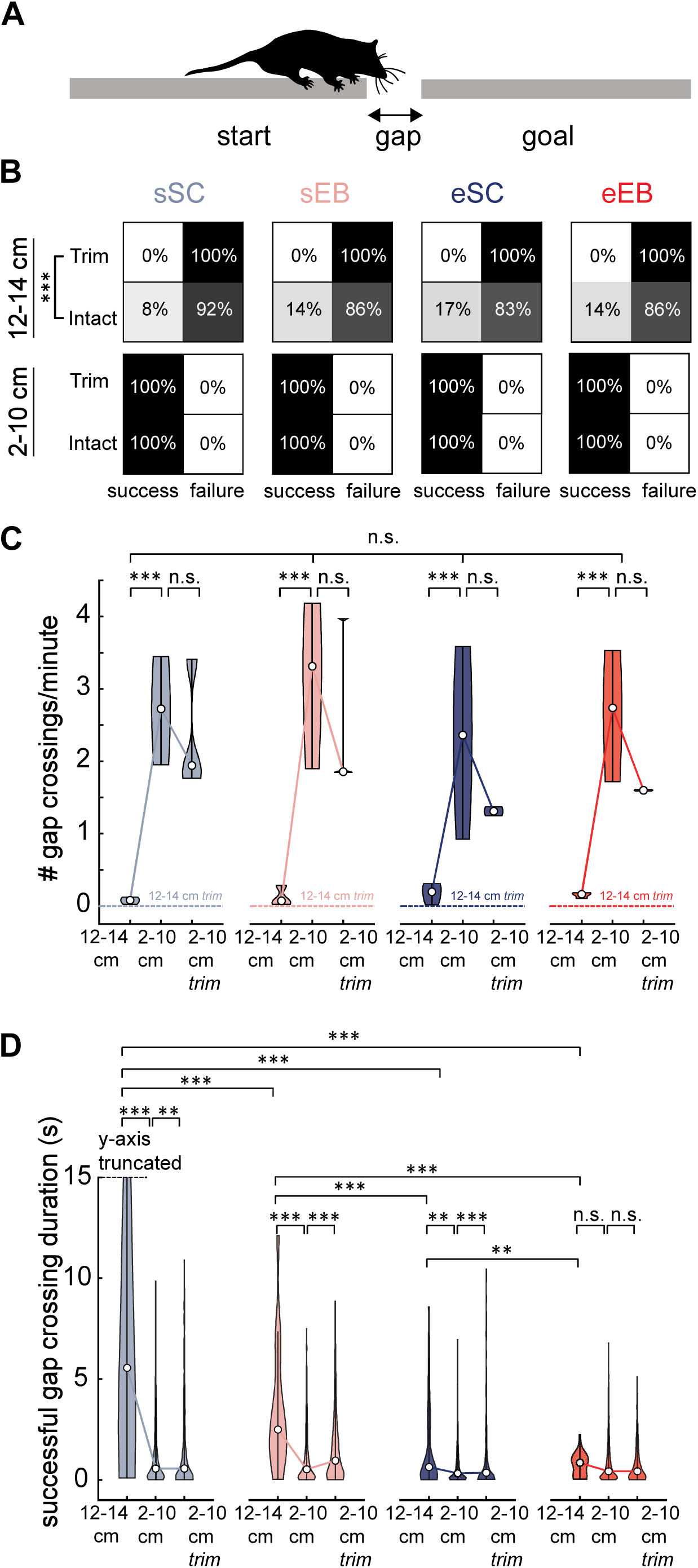
Enriched rearing alters sensorimotor performance during gap crossing. **A.** Experimental setup in the gap crossing task. **B.** Contingency tables showing the percentage of successful and failed gap crossing trials (0% = white, 100% = black; intermediate percentages shown in grayscale) for small (2–10 cm) and large (12–14 cm) gaps under intact and whisker-trimmed conditions, shown separately for each experimental group (sSC, sEB, eSC, eEB). **C.** Number of gap crossings per minute (across all trials) for large and small gaps (with intact or trimmed whiskers). Gap crossing was whisker-dependent in all groups. Data for large gaps after whisker trimming are not shown, as no successful crossings occurred in this condition. **D.** Duration of successful crossings for large and small gaps (with intact or trimmed whiskers). Most groups show significantly longer durations for succesful gap crossing on large gaps (12-14 cm) vs. small gaps (2-10 cm). Upon whisker trimming, most groups show a significant increase in gap crossing duration for small gaps, consistent with the use of whisker-dependent strategies for gap crossing. Enriched blind animals were unique in showing no significant difference between large and small gap durations and no trimming-related increase in small gap crossing times, reflecting enhanced performance on large gaps through whisker use alongside compensatory tactile strategies using other body parts that preserve performance on small gaps when whiskers are absent. For larger gaps, there are prominent differences between experimental groups based on rearing condition and blindness. Early blind animals crossed gaps faster than sighted controls in standard conditions, and their performance was further enhanced by enriched rearing. Violin plots (color-coded by experimental group) show Gaussian kernel density estimates of the data distribution, with medians indicated by circular markers. See also **Table 2**.

The fraction of successful trials (i.e. the probability of success) was strongly impacted by gap size (sSC: 8%, sEB: 14%, eSC: 17%, eEB: 14%; percent successful trials, in comparison to 100% success on small gap trials) and by whisker trimming (**Figure 3B**, Fisher’s exact test, p < 0.001; see **Table 2** for exact p-values for the fixed-effect of experimental group, and for all pairwise comparisons between groups), eliminating successful crossings at large gaps for all experimental groups. This is consistent with the role of whiskers in gap crossing described in previous studies of small mammals (Hutson & Masterton 1986, Voigts et al., 2015, Arkley et al. 2017, Graham & Socha 2020, Toso & Diamond 2023). All animals crossed on 100% of small gap trials, with or without whiskers, indicating that other body parts also contributed to small gap crossings.

Crossing rates, which reflect success rate normalized by time, were significantly lower for large gaps than for small gaps within each experimental group (**Figure 3C**, **Table 2**, **Table 3-1**; p < 0.001 in all cases, all p-values from Poisson generalized linear mixed-effects model) with no significant differences across groups (p>0.05). Although there was a trend toward decreased crossing rates in all experimental groups with whisker trimming, trimming effects were not statistically significant for crossing rates (p>0.05).

The third metric in the gap crossing task was the duration of successful crossings (**Figure 3D**, **Table 2**, **Table 3-1**). We observed significant fixed effects of experimental group (p < 0.001; all p-values from the linear mixed effects model), gap size (p < 0.001), and whisker trimming (p < 0.001), as well as interactions between group and gap size (p < 0.001) and between group and trimming (p = 0.01). Across groups, successful crossings were generally slower for large versus small gaps, with modest but significant increases in crossing times on small gaps after whisker trimming (with one exception; see below). Consistent with **Figure 3B**, this pattern suggests primary reliance on whiskers for large gaps, whereas small gaps can be crossed using strategies that incorporate both whiskers and tactile inputs from other body parts (Ramamurthy et al., 2021).

Group differences based on visual condition and rearing condition were most pronounced for large gaps (**Figure 3D**). Early blind opossums crossed large gaps faster than sighted controls under standard rearing (sSC vs. sEB: p < 0.001), and this advantage was amplified by enriched rearing (eEB vs. sEB: p < 0.001). Both enriched groups were significantly faster than standard-reared animals (p < 0.001 for all pairwise comparisons), with enriched blind animals outperforming enriched sighted animals (p < 0.01). Enriched blind animals followed the general trends of other groups but were the only cohort without a significant difference between large and small gap durations (p > 0.05), reflecting comparatively faster large gap performance. They were also the only group in which whisker trimming did not significantly prolong small gap crossings (p > 0.05). Taken together, these results indicate heightened performance supported by whisker use on large gaps, along with adaptive recruitment of other body parts to maintain small gap performance when whiskers are absent.

### Tactile experience amplifies compensatory neural plasticity in blind opossums

Single-unit recording experiments were performed in 19 opossums reared in standard conditions (10 sSC, 9 sEB), and 14 opossums reared in enriched conditions (7 eSC, 7 eEB), acquiring data for a total of 339 neurons across experimental groups (see **Table 1**; **Figure 3-1**). 63-73% of units recorded in S1 (sSC: 60, sEB: 64, eSC: 58, eEB: 51) had a significant evoked response (>2 SD above baseline) to stimulus onset following single whisker deflections, and these neurons were included in receptive field analyses. Raster plots and peristimulus time histograms (PSTHs) for whisker-responsive neurons in each group are shown in **Figure 4A-B**. These examples illustrate more selective whisker touch responses in standard-reared blind opossums, with selectivity further amplified in enriched groups.

**Figure 4.**
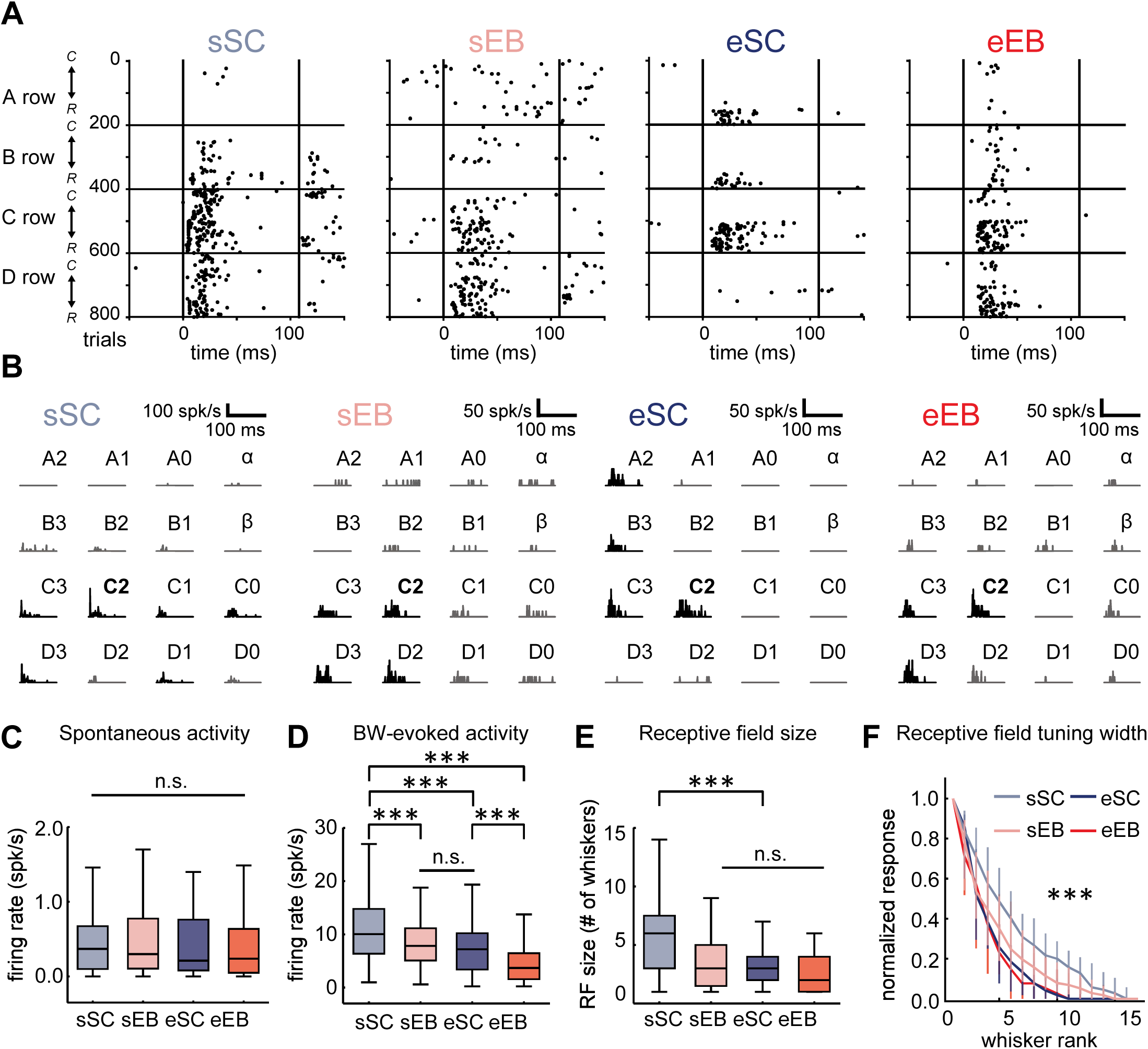
Enriched rearing amplifies alterations in S1 neuron responses to whisker touch in early blind animals and produces similar shifts in sighted controls. **A.** Spike raster plots from example neurons responsive to whisker touch in each of the four experimental groups (left to right: sSC, sEB, eSC, eEB). Examples are shown for neurons with the same best whisker (BW), C2. Trials corresponding to A, B, C, and D whisker rows are separated by horizontal lines. Within each row, the direction of more rostral (‘R’) versus more caudal (‘C’) whiskers is indicated on the y-axis. **B.** Peristimulus time histograms (PSTHs) for the same neurons shown in **(A)**. PSTHs corresponding to whiskers with response magnitudes >50% of the BW response are plotted in black, all others are in gray. The examples in **(A)** and **(B)** illustrate more selective responses to whisker touch in standard-reared blind opossums, which is further amplified in enriched groups. Additionally, neural responses to whiskers vertically adjacent to the BW are stronger than those for horizontally adjacent whiskers in enriched groups, especially in enriched early blind opossums. **C.** Spontaneous firing rates of S1 neurons were not significantly different across experimental groups. **D.** Firing rates evoked by the BW showed significant differences across experimental groups. Notably, both sighted and early blind enriched groups showed suppression relative to standard-reared sighted controls, similar to effects previously reported in early blind animals under standard rearing conditions. The magnitude of suppression of the BW-evoked response was greatest in early blind animals reared in the enriched environment. **E.** Mean receptive field size was significantly smaller in sEB, eSC and eEB animals relative to standard-reared sighted controls. This is seen both in terms of number of whiskers driving significant responses (left) and the rank-ordered tuning curve (right) for whisker-responsive S1 neurons. See also **Table 3**.

In general, spontaneous firing rates of S1 neurons were low, consistent with previously reported findings in standard-reared animals (Ramamurthy and Krubitzer, 2018), and were not significantly different among experimental groups (**Figure 4C**, **Table 3**, **Table 4-1**; mean ± SEM, sSC: 0.582 ± 0.083 spk/s, sEB: 0.613 ± 0.121 spk/s, eSC: 0.544 ± 0.079 spk/s, eEB: 0.503 ± 0.082 spk/s; linear mixed-effects model, p>0.05; see **Table 3** for exact p-values for the fixed-effect of experimental group, and for all pairwise comparisons between groups). However, responses evoked by the stimulation of even the most preferred single whisker stimulus (best whisker; BW) in enriched groups were suppressed in relation to stimulus-evoked firing rates in standard-reared sighted opossums (**Figure 4D**, **Table 3**, **Table 4-1**; mean ± SEM, sSC: 11.4 ± 0.839 spk/s, sEB: 8.39 ± 0.591 spk/s, eSC: 7.82 ± 0.748 spk/s, eEB: 4.57 ± 0.52 spk/s; linear mixed-effects model, eSC vs. sSC: p<0.01, eEB vs. sSC: p<0.001), as previously reported in early blind animals under standard rearing conditions (Ramamurthy and Krubitzer, 2018). Since relative suppression of BW-evoked responses was observed in enriched sighted controls, in addition to enriched early blind animals, this effect was not contingent on the lack of vision itself – rather it was driven by increased dependence on touch in the enriched groups of opossums. Mean BW-evoked firing rates in enriched sighted control animals were comparable to those in standard early blind animals (linear mixed-effects model, sEB vs. eSC: p>0.05). However, in enriched early blind animals, mean BW-evoked firing rates were lower relative to all other experimental groups (linear mixed-effects model, eEB vs. sEB: p<0.001, eEB vs. eSC: p<0.001, eEB vs. sSC: p<0.001). Thus, blindness was not required for suppression of whisker-evoked responses, but the combination of tactile enrichment and vision loss produced the greatest effect on whisker-evoked firing rates of S1 neurons. This is consistent with animals relying on their whiskers to a greater extent when reared in a complex environment in the absence of vision.

Average whisker receptive fields in both sighted and blind enriched groups were significantly smaller than those in standard-reared sighted controls (mean ± SEM, eEB: 3.22 ± 0.409, eSC: 3.48 ± 0.364, sSC: 5.92 ± 0.427; linear mixed-effects model, eSC vs. sSC: p<0.001, eEB vs. sSC: p<0.001) when measured as the number of whiskers that evoked a significant response above spontaneous firing (**Figure 4E**, left, **Table 3**, **Table 4-1**). We see similar results when receptive fields are quantified as the mean rank-ordered tuning curve across neurons (**Figure 4E**, right, **Table 3**, **Table 4-1**; linear mixed-effects model, eSC vs. sSC: p<0.001). The finding that tactile enrichment in sighted control animals leads to a decrease in mean receptive field size—similar to both standard-reared and enriched early blind animals—supports the interpretation that this reduction reflects an experience-dependent effect across all three groups. However, changes in measures of overall receptive field size in enriched early blind animals relative to standard-reared early blind animals were either not significant (**Figure 4E**, left, number of whiskers: eEB: 3.22 ± 0.409, sEB: 3.38 ± 0.270; linear mixed-effects model, eEB vs. sEB: p>0.05) or modest (**Figure 4E**, right, rank-ordered tuning curve: linear mixed-effects model, p<0.01). This suggests that, under standard rearing conditions, receptive field plasticity in early blind animals may already have reached a limit that cannot be further shifted by enrichment.

### Enrichment enhances selectivity for whisker touch along the behaviorally relevant horizontal axis

To better understand the effects of enriched rearing on receptive field configuration of S1 neurons, we characterized two-dimensional receptive fields for the row (horizontal) and arc (vertical) dimensions of the whisker pad (**Figure 5A**). Visual inspection of mean 2D receptive fields revealed clear differences in receptive field anisotropies across experimental groups (**Figure 5B**). We quantified anisotropies in receptive fields as the *shape index*, ranging in value from −1 to 1, where positive values indicated horizontally anisotropic shapes, negative values indicated vertically anisotropic shapes and values close to 0 indicated more isotropic shapes. Receptive fields for neurons in the S1 whisker representation in standard-reared sighted control and early blind animals were dominated by horizontally anisotropic shapes (**Figure 5C-D**, **Table 3**, **Table 5-1**). However, for early blind animals reared in enriched conditions, the receptive field shape distributions were shifted away from horizontal anisotropy – either when considering only 1° surround whisker (SW) positions (**Figure 5C**; neurons with shape index > 0.0, sSC: 68.3%, sEB: 57.8%, eEB: 37.3%, Fisher’s exact test, eEB vs. sSC: p=0.001, eEB vs. sEB: p<0.05; linear mixed-effects model, eEB vs. sSC: p<0.001, eEB vs. sEB: p<0.05), or when including all SW (1° - 3°) positions (**Figure 5D**; neurons with shape index > 0.0, eEB: 37.3%, sSC: 81.7%, sEB: 76.6%, Fisher’s exact test, eEB vs. sSC: p<0.001, eEB vs. sEB: p<0.001; linear mixed-effects model, eEB vs. sSC: p<0.001, eEB vs. sEB: p<0.001).

**Figure 5.**
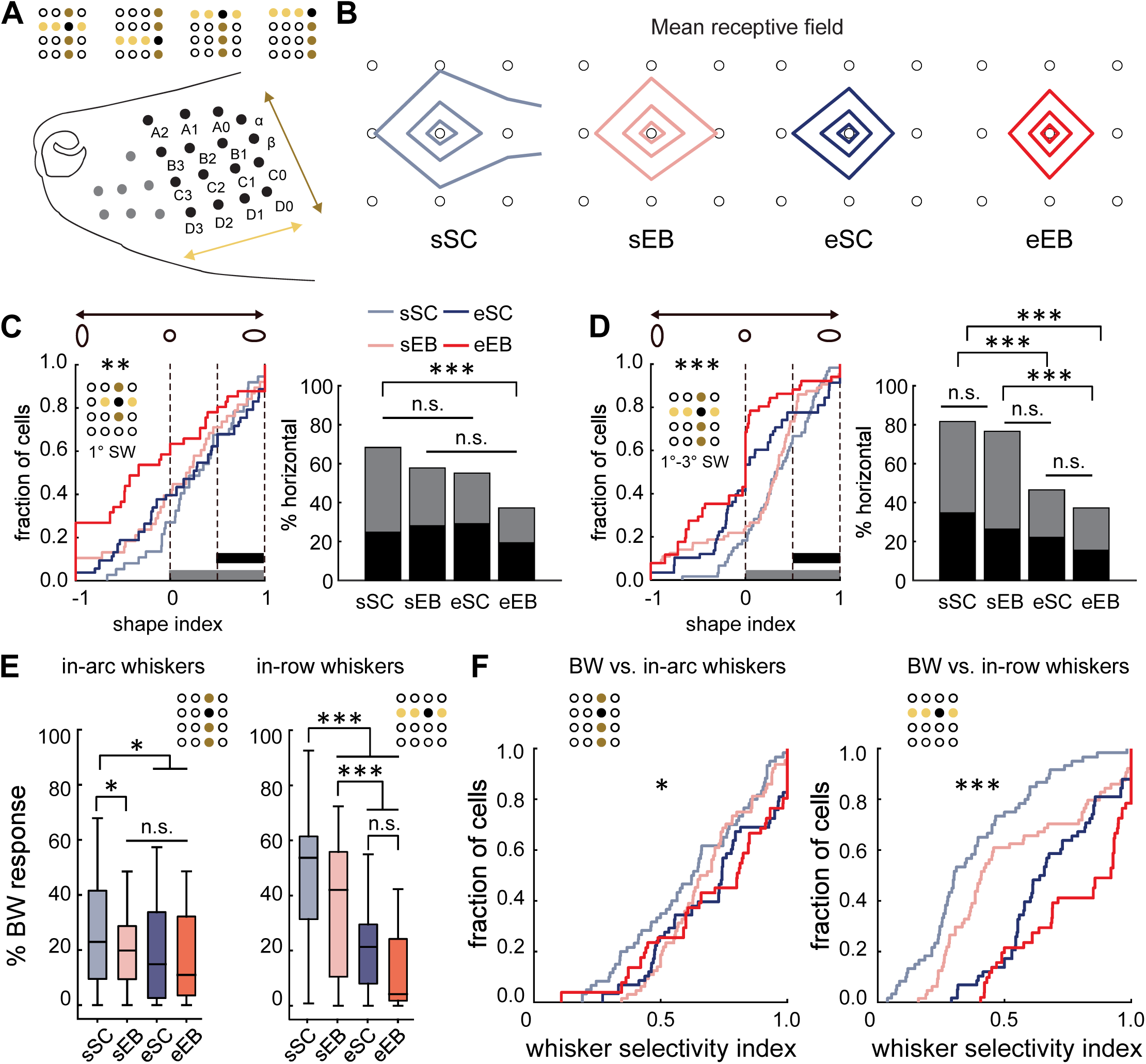
Enriched rearing amplifies anisotropies in receptive fields of S1 neurons in early blind animals, improving selectivity for whisker touch along a behaviorally relevant axis. **A.** Schematic of the opossum mystacial whisker pad indicating whisker rows (light yellow) and arcs (gold), and the horizontal and vertical axes used for receptive field shape analyses. Insets in individual panels depict the grid of 16 whiskers and the surrounding whisker (SW) positions included in each analysis, shown for one example best whisker (BW) position (black filled circle). **B.** Mean two-dimensional somatotopic receptive fields for each experimental group (left to right: sSC, sEB, eSC, eEB), shown with 50%, 75%, and 90% contour lines (response levels relative to the BW, smoothed by linear interpolation for display). This visualization highlights the degree of horizontal vs. vertical spread (anisotropy) in the average receptive field configuration for each experimental group. **C-D.** Receptive field shape distributions for S1 neurons in enriched animals were shifted away from horizontal anisotropic shapes, which are typical of standard-reared animals. **C** shows receptive field shape distributions when only 1° SW positions were considered (left) and fractions of horizontally anisotropic receptive fields corresponding to each group (right). Light gray bars indicate horizontally elongated receptive fields defined as shape indices > 0, while black bars indicate horizontally elongated receptive fields with a stricter cutoff of shape indices > 0.5. **D** shows data for the same neurons when all SW positions (1-3°) were included. Horizontally anisotropic receptive fields are decreased in both enriched groups of animals, and to the greatest extent in enriched early blind animals. **E.** Quantification of tuning width along the row (horizontal/rostrocaudal axis) and arc (vertical axis) across experimental groups. Enriched groups show sharper tuning along the horizontal axis, to a significantly greater extent than standard-reared early blind animals. Enriched groups also show sharper tuning along the vertical axis compared with standard-reared sighted controls, but only to the same extent as previously reported in standard-reared early blind animals. **F.** Distributions of the whisker selectivity index of S1 neurons calculated separately for in-arc whiskers (left) and in-row whiskers (right), when including all SW positions (1°-3°). sEB, eSC and eEB groups all showed significantly greater selectivity for whiskers within arcs, but were not significantly different from each other. The greatest increases in whisker response selectivity were seen for whiskers within rows i.e. the horizontal/rostrocaudal axis, which is the primary axis of natural whisker motion. Selectivity was greatest for eEB animals followed by eSC and sEB animals, with all three of these groups showing significantly higher selectivity along the horizontal axis compared to sSC animals. See also **Table 3**.

Receptive field distributions were also shifted away from horizontal anisotropy in sighted control animals reared in the enriched environment. This shift was not seen when only 1° SW positions were considered (**Figure 5C**; neurons with shape index > 0.0, eSC: 55.2%, Fisher’s exact test, eSC vs. sSC: p>0.05, eSC vs. sEB: p>0.05; linear mixed-effects model, eSC vs. sSC: p>0.05, eSC vs. sEB: p>0.05) but becomes apparent when 1° - 3° SW positions are considered (**Figure 5D**; neurons with shape index > 0.0, eSC: 46.6%, Fisher’s exact test, eSC vs. sSC: p<0.001, eSC vs. sEB: p<0.001; linear mixed-effects model, eSC vs. sSC: p=0.01, eSC vs. sEB: p>0.05). Thus, rearing in an enriched environment, which was permissive of more naturalistic patterns of tactile behavior, shifted the shapes of receptive fields of S1 neurons away from the horizontally anisotropic shapes typically found in animals reared in standard laboratory cages. The shift occurred in both sighted and early blind enriched animals, but the effect was more pronounced in early blind animals.

The results of receptive field shape analyses pointed to differential effects of rearing environment on tuning along horizontal and vertical axes of receptive fields. We quantified neuronal tuning width separately along whisker rows and arcs for each experimental group (**Figure 5E**, **Table 3**, **Table 5-1**). Both sighted and early blind enriched animals showed sharper tuning along both the vertical and horizontal axes relative to sighted controls, but there was a striking difference in the magnitude of tuning width changes along the horizontal vs. vertical axes.

This constitutes an amplification of shifts in receptive field anisotropies previously described in early blind animals under standard rearing conditions (Ramamurthy and Krubitzer, 2018). Along the arc (vertical) axis, tuning was sharpened to the same extent in the enriched groups as seen in standard-reared early blind animals relative to sighted controls (**Figure 5E**; %BW response for in-arc whiskers: eEB: 19.4 ± 2.80%, eSC: 18.7 ± 2.13%, sEB: 20.1 ± 1.58%, sSC: 26.7 ± 2.50%; linear mixed-effects model, eEB vs. sSC: p<0.05, eEB vs. sEB: p>0.05, eSC vs. sSC: p<0.05, eSC vs. sEB: p>0.05). But along the row (horizontal axis), both sighted and early blind enriched animals showed sharper tuning relative to both groups of standard-reared animals (**Figure 5E**; %BW response for in-row whiskers: eEB: 13.8 ± 2.07%, eSC: 20.9 ± 1.96%, sEB: 35.1 ± 2.82%, sSC: 49.1 ± 2.85%; linear mixed-effects model, eEB vs. sSC: p<0.001, eEB vs. sEB: p<0.001, eSC vs. sSC: p<0.001, eSC vs. sEB: p<0.001). There was a trend for sharper row tuning in enriched early blind animals compared to enriched sighted control animals, but this was not significant (linear mixed-effects model; eSC vs. eEB: p>0.05). We computed the whisker selectivity index as the normalized difference ratio of the neuronal response to the best whisker in comparison to responses to in-row or in-arc whiskers. Population distributions of the selectivity index for S1 neurons (**Figure 5F**, **Table 3**, **Table 5-1**) further validate the anisotropy in the enhancement of selectivity seen for animals reared in enriched environments (BW vs. in-arc whiskers: linear mixed-effects model, eEB vs. sSC: p<0.05, eEB vs. sEB: p>0.05, eSC vs. sSC: p<0.05, eSC vs. sEB: p>0.05, eEB vs. eSC: p>0.05; BW vs. in-row whiskers: linear mixed-effects model, eEB vs. sSC: p<0.001, eEB vs. sEB: p<0.001, eSC vs. sSC: p<0.001, eSC vs. sEB: p<0.001, eEB vs. eSC: p>0.05).

Thus, a spatially complex enriched environment drives significantly greater improvement in neuronal selectivity for whisker touch in early blind animals especially along the rostrocaudal axis, which is the primary axis of natural whisking behavior. A similar improvement in selectivity was observed in sighted control littermates reared in the enriched environment, supporting the role of tactile experience rather than vision loss itself in driving these effects. In summary, the results of our current study indicate that compensatory effects on neural coding in S1 following early blindness do not arise solely from visual deprivation and can be amplified and directed by tactile experience.

## DISCUSSION

Although the effects of enrichment on plasticity within a sensory modality have been studied at structural, functional, and molecular levels, few studies have examined its impact on cross-modal compensatory plasticity, focusing mainly on c-fos expression or synaptic changes (Piche et al., 2004; Mundiñano and Martínez-Millán, 2010; Zheng et al., 2014). The short-tailed opossum is born with a highly immature nervous system, enabling precise extrauterine manipulations at very early stages in development (Ramamurthy and Krubitzer, 2018; Ramamurthy et al., 2021). Here, we tested the effects of rearing sighted and early blind short-tailed opossums in a large and dynamic, spatially complex 3D environment, and examined effects on tactile behavior and receptive field configurations of neurons in S1.

Enrichment reduces abnormal stereotypic behaviors (e.g. overgrooming) and other behavioral indicators of stress and aggression, and enhances learning, memory, motor performance and social behavior (Dijkhuizen et al., 2024; Ratuski et al., 2024; Cait et al., 2024; Nakhwa et al., 2024; Caires & Bossolani-Martins, 2023; Mieske et al., 2022; Kentner et al., 2021; Khoo et al., 2020; Bailoo et al., 2018; Girbovan & Plamondon, 2013; Amaral et al., 2008; Mason et al., 2007). It can also improve sensory function and performance, including whisker touch discrimination (Zheng et al., 2021). In our study, we implemented enrichment intended to increase the use of whisker touch. Since whiskers are critical for guiding forelimb placement in crossing gaps when small mammals climb (Ahl, 1986; Arkley et al., 2014; Niederschuh et al., 2015; Arkley et al., 2017; Grant et al., 2018), we reared opossums in cages with periodically varying, spatially complex climbing substrates and intermittently introduced novel toys, since novel objects and other interactive enrichment devices promote active whisking behavior (Nakhwa et al., 2024; Sofroniew & Svoboda, 2015). We show that in a novel setting, both sighted and blind opossums showed differential exploration patterns reflecting the geometry of their rearing environments. Additionally, enriched opossums are faster and more successful in gap crossing using a whisker-dependent strategy. Thus, opossums use behavioral strategies shaped by their rearing environments, and our enrichment approach effectively engages and enhances naturalistic whisker-mediated behavior in both sighted and blind animals, enabling us to evaluate the effects of enriched rearing on compensatory plasticity following early vision loss.

It has been previously reported that early blindness alters the coding of whisker touch inputs in opossums, in a manner that supports enhanced tactile discrimination (Ramamurthy and Krubitzer, 2018; Ramamurthy et al., 2021). This involved suppression of whisker-evoked responses combined with increased spatial selectivity, due to the greater relative suppression of neural responses to non-preferred stimuli (SW) vs. the preferred stimulus (BW). These alterations in the receptive fields of neurons in S1 resemble the effects of naturalistic environmental enrichment on receptive fields of neurons in S1 of adult rats (Polley et al., 1999, 2004). Here we demonstrate that rearing early blind short-tailed opossums in an enriched environment further amplifies these changes. Since selectivity to whisker stimuli was enhanced not just in early blind animals but also in sighted control animals reared in enriched conditions, we can infer that suppression of whisker-evoked responses and increased selectivity of neural responses in the S1 whisker representation are effects of tactile experience rather than visual deprivation itself.

A key finding was the effect of the rearing environment on the shapes of receptive fields of S1 neurons – shape distributions for both sighted and early blind animals exhibited a prominent shift away from the horizontally anisotropic shapes that dominate in standard-reared opossums. This was accompanied by a reduction in tuning width along row (horizontal) vs. arc (vertical) axes in enriched animals – on average, tuning widths were reduced to a greater extent along the row axis, parallel to the main axis of whisker movements. This may reflect more naturalistic patterns of whisker use in short-tailed opossums, enabled by the increase in vertical space and availability of climbing substrates in the enriched rearing environment. In rats, simulations that modelled exploratory behavior predicted changes in the spatial patterns of whisker contact depending on the environment being explored (Hobbs et al., 2015) – whiskers within the same horizontal row were more likely to come in simultaneous contact during exploration of flat surfaces, but in more complex and naturalistic environments, the probability distributions of simultaneous whisker contact shifted away from this row-wise structure. Such alterations in the patterns of tactile stimuli impinging on the whisker array over the course of development may, in turn, impact receptive field structure.

Another possibility is that anisotropic effects on receptive fields are inherent to differences in row vs. arc organization within the neocortex. In rats and mice, it is well-documented that there is a bias towards preferential connectivity among barrels corresponding to whiskers in the same row (Bernardo et al., 1990; Hoeflinger et al., 1995; Kim and Ebner, 1999; Keller and Carlson, 1999), and neurons in barrel cortex typically display patterns of activity that tend to be elongated along the row axis (Armstrong-James and Fox, 1987; Simons, 1978; Kleinfeld and Delaney, 1996; Ego-Stengel et al., 2005; Le Cam et al., 2011). Further, there is evidence for differential effects on responses of S1 neurons along row and arc axes when sensory inputs are altered within the whisker system, for example, through chronic stimulation (Quairiaux et al., 2007) or selective trimming (Dubroff et al., 2005; Wallace and Sakmann, 2008) of specific whiskers. Thus, biases in the functional organization and connectivity within the S1 whisker representation may favor plasticity, preferentially sharpening tuning along the row axis.

A long-standing question has been whether reorganization following sensory loss is primarily driven by deprivation of the lost modality or due to increased experience using the spared senses. Studies in blind humans with Braille reading experience support an important role for tactile experience in generating tactile spatial acuity enhancement following blindness (Wong et al., 2011). The relative contributions of deprivation of the lost modality versus increased or modified use of the spared senses (Goldreich and Kanics, 2003; Wong et al., 2011; Voss, 2011) to different aspects of cortical reorganization following sensory loss are still not well understood. Our study provides clear evidence for the role of experience in driving compensatory neural and behavioral plasticity following sensory loss, highlighting the importance of therapeutic strategies engaging tactile exploration (for example through play-based learning of tactile skills, Hilditch 2024; or the use of haptic devices, Fleck et al., 2025) to maximize neural and behavioral compensation through touch.

## Acknowledgments

We thank Dr. Cindy Clayton for animal care, Carly Jones for assistance with preliminary data analysis, and Carlos Pineda for comments on a draft of this manuscript.

## MULTIMEDIA

**Movie 1.** A nesting mother opossum navigating the enriched home cage environment. Movie shown at 4x speed.

**Movie 2.** Sighted opossum in the arena test. Movie shown at 4x speed. **Movie 3.** Early blind opossum in the arena test. Movie shown at 4x speed. for Neuroscience 34:5406-5415.

## EXTENDED DATA FIGURE LEGENDS

**Figure 1-1. Standard and enriched rearing environments used for short-tailed opossums. A.** Comparison of cages used in standard (left) and enriched (right) rearing paradigms. Enriched cages (29”l x 18”w x 24”h) were substantially larger (∼8x in volume) than standard cages (8.5”l x 10”w x 8”h), especially in the vertical dimension (∼3x). Each enriched cage contained a nesting box positioned high above the ground, in addition to the standard nesting cup on the cage floor. Cages also included various enrichment objects: regularly repositioned manzanita branches, rotating sets of enrichment toys, a running wheel, and social housing (see *Materials and Methods*). **B.** Frontal view of an enriched cage. **C.** Top view of the same enriched cage shown in **(B).** Letters mark individual components visible in both views: nesting box (a), nesting cup (b), food (c), water (d), running wheel (e), manzanita branches (f–l), and enrichment toys (m). **D.** Mother with an attached experimental litter after placement in the enriched rearing environment. **E–F.** Early blind weanling climbing up a branch to reach the nesting box.

**Figure 2-1. Behavioral movie analyses. A.** Representative frame showing semi-automated identification of an opossum during arena testing using idtracker.ai. **B.** Representative frame showing behavioral tracking in the gap crossing task using DeepLabCut. “IR1” and “IR2” indicate the positions of infrared motion sensors mounted on the edge of each platform, tracked with DeepLabCut to aid in gap crossing analyses. **C.** Zoomed-in view showing an opossum on the start platform extending its whiskers toward the goal platform prior to crossing the gap.

**Figure 3-1. Histological verification of recording sites. A, C, E, G.** Myelin-stained tangential sections of cortex from a standard-reared sighted (sSC) animal **(A)**, a standard-reared early blind (sEB) animal **(C)**, an enriched sighted (eSC) animal **(E)**, and an enriched early blind (eEB) animal **(G)**. Primary somatosensory cortex (S1) is identifiable as a darkly staining region relative to adjacent cortical fields. **B, D, F, H.** Reconstructions of myeloarchitectural borders drawn from the series of myelin-stained sections in sSC (**B**), sEB (**D**), eSC (**F**) and eEB (**H**) animals. The locations of fluorescent probe insertions are shown as open circles, and recording sites are shown as black filled circles. Only sites at which whisker-evoked neuronal responses were quantified are included. In all panels, medial (M) and rostral (R) directions are indicated by arrows. Scale bar = 250 µm.

## EXTENDED DATA TABLES

**Table 3-1. ANOVA marginal tests for fixed effects in Figure 3**. F statistics, numerator and denominator degrees of freedom (DF1, DF2), and p-values are reported for each model term in analyses corresponding to **Figure 3**.

**Table 4-1. ANOVA marginal tests for fixed effects in Figure 4**. F statistics, numerator and denominator degrees of freedom (DF1, DF2), and p-values are reported for each model term in analyses corresponding to **Figure 4**.

**Table 5-1. ANOVA marginal tests for fixed effects in Figure 5**. F statistics, numerator and denominator degrees of freedom (DF1, DF2), and p-values are reported for each model term in analyses corresponding to **Figure 5**.

