## Extended Data for "Complex three-dimensional rearing environments amplify compensatory plasticity following early blindness"

Figure 1-1

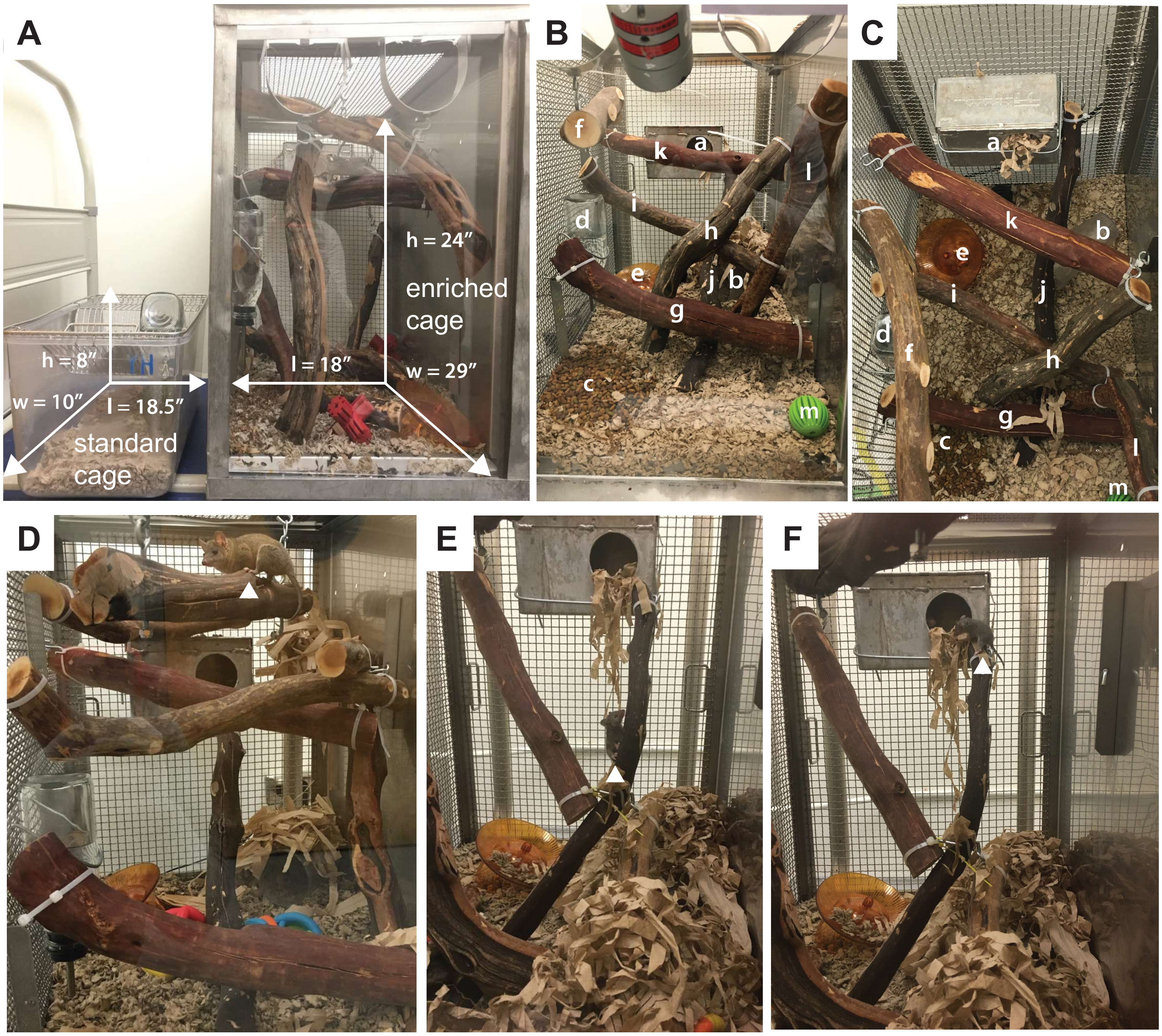
A
h = 8″
w = 10″
l = 18.5″
standard cage
h = 24″
enriched cage
l = 18″
w = 29″
B
f
k
a
l
i
d
h
e
j
b
g
c
m
C
a
k
b
e
i
j
d
f
h
g
c
l
m
D
E
F

Figure 2-1

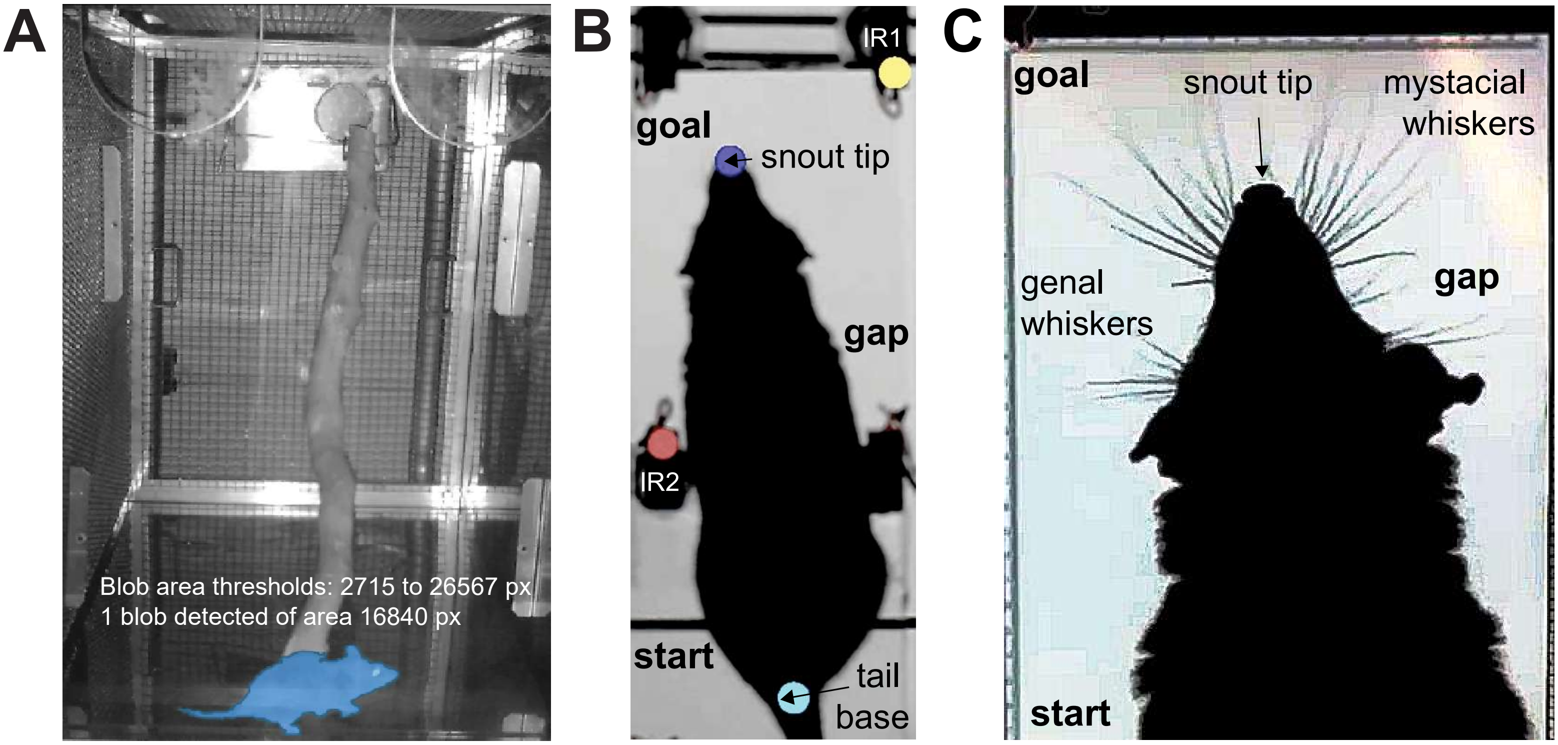

A
Blob area thresholds: 2715 to 26567 px
1 blob detected of area 16840 px
B
IR1
goal
snout tip
gap
IR2
start
tail
base
C
goal
snout tip
mystacial
whiskers
genal
whiskers
gap
start

Figure 3-1

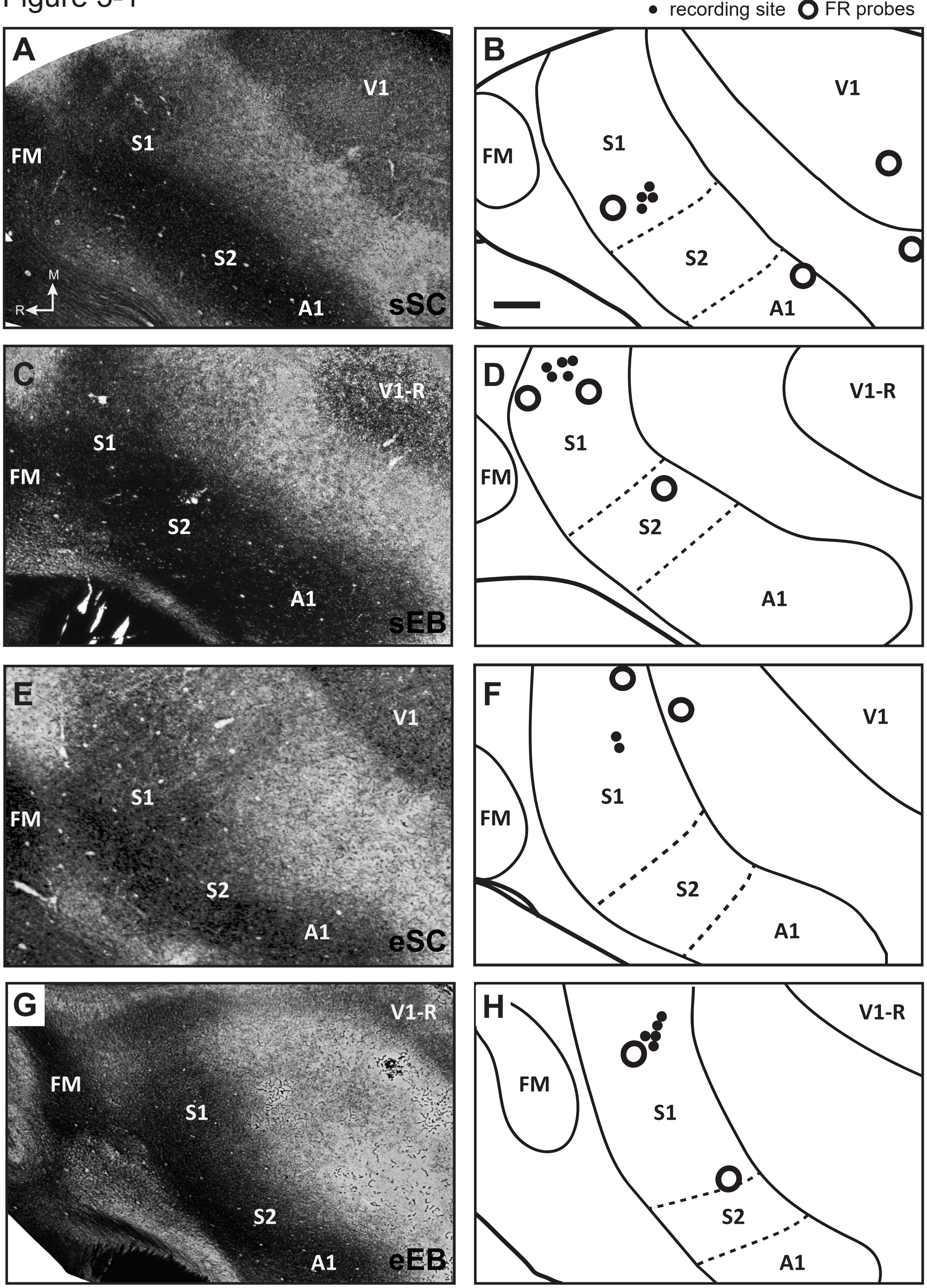

recording site
FR probes
A
V1
S1
FM
S2
A1
M
R
sSC
B
V1
S1
FM
S2
A1
C
V1-R
S1
FM
S2
A1
sEB
D
V1-R
S1
FM
S2
A1
E
V1
S1
FM
S2
A1
eSC
F
V1
S1
FM
S2
A1
G
V1-R
FM
S1
S2
A1
eEB
H
V1-R
FM
S1
S2
A1

### EXTENDED DATA TABLES

#### Extended Data Table 3-1. ANOVA marginal tests for fixed effects in Figure 3.

| Figure | ANOVA marginal tests | FStat | DF1 | DF2 | pValue |
|---|---|---|---|---|---|
| **Figure 3C** | **ANOVA marginal tests: DFMethod = 'residual'** | | | | |
| | Term | FStat | DF1 | DF2 | pValue |
| | {'(Intercept)'} | 21.367 | 1 | 24 | 0.00010846 |
| | {'ExptGp' } | 0.64661 | 3 | 24 | 0.5927 |
| | {'Gap' } | 84.689 | 1 | 24 | 2.4277e-09 |
| | {'Trim' } | 0.39504 | 1 | 24 | 0.5356 |
| | {'ExptGp:Gap' } | 1.6189 | 3 | 24 | 0.21123 |
| | {'ExptGp:Trim'} | 0.16617 | 3 | 24 | 0.91812 |
| **Figure 3D** | **ANOVA marginal tests: DFMethod = 'Satterthwaite'** | | | | |
| | Term | FStat | DF1 | DF2 | pValue |
| | {'(Intercept)'} | 678.83 | 1 | 10514 | 4.539e-145 |
| | {'Gap' } | 421.27 | 1 | 10514 | 8.0367e-92 |
| | {'Trim' } | 7.013 | 1 | 10514 | 0.0081039 |
| | {'ExptGp' } | 23.687 | 3 | 10514 | 2.8406e-15 |
| | {'Gap:ExptGp' } | 111.98 | 3 | 10514 | 2.2426e-71 |
| | {'Trim:ExptGp'} | 9.3432 | 3 | 10514 | 3.6434e-06 |

F statistics, numerator and denominator degrees of freedom (DF1, DF2), and p-values are reported for each model term in analyses corresponding to **Figure 3**.

#### Extended Data Table 4-1. ANOVA marginal tests for fixed effects in Figure 4.

| Figure | ANOVA marginal tests | FStat | DF1 | DF2 | pValue |
|---|---|---|---|---|---|
| **Figure 4C** | **ANOVA marginal tests: DFMethod = 'Satterthwaite'** | | | | |
| | Term | FStat | DF1 | DF2 | pValue |
| | {'(Intercept)'} | 39.173 | 1 | 339 | 1.1718e-09 |
| | {'ExptGp' } | 0.25773 | 3 | 339 | 0.85581 |
| **Figure 4D** | **ANOVA marginal tests: DFMethod = 'Satterthwaite'** | | | | |
| | Term | FStat | DF1 | DF2 | pValue |
| | {'(Intercept)'} | 140.79 | 1 | 4.7301 | 0.00010652 |
| | {'ExptGp' } | 17.101 | 3 | 232.42 | 4.5371e-10 |
| **Figure 4E (right)** | **ANOVA marginal tests: DFMethod = 'Satterthwaite'** | | | | |
| | Term | FStat | DF1 | DF2 | pValue |
| | {'(Intercept)'} | 266.75 | 1 | 233 | 1.7587e-40 |
| | {'ExptGp' } | 12.369 | 3 | 233 | 1.5486e-07 |
| **Figure 4E (left)** | **ANOVA marginal tests: DFMethod = 'Satterthwaite'** | | | | |
| | Term | FStat | DF1 | DF2 | pValue |
| | {'(Intercept)'} | 1188.9 | 1 | 3483 | 2.0524e-224 |
| | {'ExptGp' } | 40.206 | 3 | 3483 | 1.5495e-25 |

F statistics, numerator and denominator degrees of freedom (DF1, DF2), and p-values are reported for each model term in analyses corresponding to **Figure 4**.

**Extended Data Table 5-1. ANOVA marginal tests for fixed effects in Figure 5.**

| | ANOVA marginal tests: DFMethod = 'Satterthwaite' | | | | |
|---|---|---|---|---|---|
| **Figure 5C** | Term | FStat | DF1 | DF2 | pValue |
| | {'(Intercept)'} | 8.7766 | 1 | 169 | 0.0034913 |
| | {'ExptGp' } | 4.6638 | 3 | 169 | 0.0037027 |
| **Figure 5D** | **ANOVA marginal tests: DFMethod = 'Satterthwaite'** | | | | |
| | Term | FStat | DF1 | DF2 | pValue |
| | {'(Intercept)'} | 27.068 | 1 | 233 | 4.3037e-07 |
| | {'ExptGp' } | 8.1681 | 3 | 233 | 3.4152e-05 |
| **Figure 5E (left)** | **ANOVA marginal tests: DFMethod = 'Satterthwaite'** | | | | |
| | Term | FStat | DF1 | DF2 | pValue |
| | {'(Intercept)'} | 146.11 | 1 | 233 | 1.9597e-26 |
| | {'ExptGp' } | 2.7797 | 3 | 233 | 0.041847 |
| **Figure 5E (right)** | **ANOVA marginal tests: DFMethod = 'Satterthwaite'** | | | | |
| | Term | FStat | DF1 | DF2 | pValue |
| | {'(Intercept)'} | 391.43 | 1 | 233 | 8.7398e-52 |
| | {'ExptGp' } | 37.602 | 3 | 233 | 7.4223e-20 |
| **Figure 5F (left)** | **ANOVA marginal tests: DFMethod = 'Satterthwaite'** | | | | |
| | Term | FStat | DF1 | DF2 | pValue |
| | {'(Intercept)'} | 156.86 | 1 | 233 | 7.4003e-28 |
| | {'ExptGp' } | 34.262 | 3 | 233 | 2.2108e-18 |
| **Figure 5F (right)** | **ANOVA marginal tests: DFMethod = 'Satterthwaite'** | | | | |
| | Term | FStat | DF1 | DF2 | pValue |
| | {'(Intercept)'} | 461.91 | 1 | 233 | 3.2951e-57 |
| | {'ExptGp' } | 2.7633 | 3 | 233 | 0.042756 |

F statistics, numerator and denominator degrees of freedom (DF1, DF2), and p-values are reported for each model term in analyses corresponding to **Figure 5**.
